# Friend, not Foe: Lowered Tissue Reactivity to Long-Term Polyimide Implants

**DOI:** 10.64898/2026.02.06.703281

**Authors:** Corinne Orlemann, Laura M. De Santis, Paul Neering, Christian Boehler, Kirti Sharma, Arno Aarts, Tobias Holzhammer, Rik J.J. van Daal, Patrick Ruther, Maria Asplund, Roxana N. Kooijmans, Pieter R. Roelfsema

## Abstract

One of the biggest challenges for neurotechnology is the design of devices that are tolerated well by brain tissue, without sacrificing functionality and implantability. This study examined which design choices mitigate tissue damage and improve longevity, by varying probe features implanted in the cerebral cortex of mice.

We report on a systematic, quantitative analysis of neuronal and inflammation markers across cortical depth. We implanted a total of 103 stiff silicon or flexible polyimide probes in 32 mice, varying their thicknesses and widths, which were either attached to the skull or not. A new, automated workflow to quantify immunohistochemical data examines: 1) the tissue loss caused by the implant, 2) the cortical neuronal density, and 3) the immune response expressed by astrocytic and microglial reaction.

Flexible polyimide probes exhibited a clear advantage, with fewer lesions and weaker immune responses than stiff silicon probes. Furthermore, we observed a weaker influence of the shank cross-section. A cortical depth profile of immune reactivity revealed focal reactions at the device entry points in the superficial cortex and at the cortex-white matter boundary. This study gives important insights on optimizing device design parameters as well as surgical insights for improved tissue integration of intracortical electrode arrays.

## Introduction

Over the past years the field of neurotechnology has seen a rapid advancement of technological developments for interfacing with the brain(Dalrymple et al., 2025; Vázquez-Guardado et al., 2020). One of the most reliable strategies is to implant microelectrode arrays into the brain, which can record brain signals at a high spatial and temporal resolution and stimulate neurons by delivering weak electrical currents. These neuronal probes have been successfully applied to read and write to the brain, retrieving motor commands from the brain of paralyzed individuals (Collinger et al., 2013; Hochberg et al., 2012) and generate sensations in prosthetic applications for touch and vision (Chen et al., 2020; Fernández et al., 2021; Flesher et al., 2016).

A main limitation of intracortical neuronal probes is their adverse effect on the targeted neuronal networks (Biran et al., 2007; Otte et al., 2022; Wellman et al., 2019; Woolley et al., 2013). Implantation requires advancing the probe through an opening in the skull and into the brain tissue, inherently damaging neurons and blood vessels along its path (Kook et al., 2016; Otte et al., 2022). The tissue reaction to an implanted foreign body is twofold (McConnell et al., 2009; Polikov et al., 2005; Szarowski et al., 2003). During the implantation, the probe ruptures the blood vessels and cuts through neuronal connections, triggering an initial acute inflammatory reaction. This reaction primarily consists of activated microglial cells responding to the injury and lasts about one to two weeks (Kozai et al., 2012; Potter et al., 2012). After this primary tissue response, a second, chronic inflammatory response occurs. This chronic reaction is characterized by the formation of scar tissue with reactive astrocytes along the electrode track and the loss of neurons around the site of implantation (Salatino et al., 2017). So far, the only commercially available devices are rigid silicon probes, like the Utah-array (Campbell et al., 1991; Obidin et al., 2020) and more recently, active stiff devices such as the Neuropixels (Steinmetz et al., 2021), the SiNAPS probe (Angotzi et al., 2019) or fully immersible probes as introduced by De Dorigo et al. (2018). Active devices minimize the space occupied by interconnection lines, substantially reducing the probe cross-section. Several studies report on the ability to record neuronal activity with these devices over months to years, but there is an invariable decline in device functionality over time (Barrese et al., 2013). The glial scarring and neuronal damage caused by the chronic inflammation is one of the major reasons quoted for the loss of functionality (Polikov et al., 2005; Szarowski et al., 2003). Reactive astrocytes form a sheath around the implant and replace the neurons near the electrodes, increasing the distance of the microelectrodes to the closest neurons they can record or stimulate. This adverse response decreases recording quality and increases currents necessary for electrical stimulation (Chen et al., 2023). In recent years, researchers and device manufacturers started to shift towards the use of highly flexible polymer-based probes (Du et al., 2017; Lind et al., 2013; Luan et al., 2017; Nguyen et al., 2014; Park et al., 2021; Tian et al., 2023; Zhao et al., 2023), aiming to counteract the adverse effects of rigid silicon-based probes on the brain tissue. Due to their flexibility, these probes can follow the movements of the brain tissue, and the material allows the fabrication of thinner probes that do not break when implanted into the brain. Several studies have reported high functionality of such devices, which can record brain signals over a long period of time and stimulate neurons with low currents (Chung et al., 2019; Lycke et al., 2023; Orlemann et al., 2024; Yasar et al., 2024).

Previous studies examined how the size of the probes and the surgical implantation strategy influence how well they integrate in the brain tissue. Bulkier probes replace more tissue and cause more neuronal damage than probes of smaller size (Seymour & Kipke, 2007; Stice et al., 2007; Thelin et al., 2011), and in case of flexible probes, thicker probes are more rigid than thinner ones. Furthermore, silicon probes can be pushed into the brain because they are rigid, but polyimide probes are most often inserted using shuttle devices, which are retracted upon implantation so that only the flexible probe stays behind. The insertion of these shuttle devices inevitably increases the implantation footprint, causing additional tissue loss (Otte et al., 2022). The connection between the probe and the connector plays another important factor in the development of neural devices. Many current devices are tethered; the brain implant is connected to a fixed position, e.g. a printed circuit board (PCB) on the skull. Prior studies have shown that this tethering can induce additional chronic trauma to the brain, presumably because it limits the ability of the probe to follow the micromovements of the brain (Biran et al., 2007; Kim et al., 2004; Thelin et al., 2011; Vomero et al., 2022).

Despite rapid advancements in neurotechnology, it is still uncertain which design parameters play the most important role in promoting tissue integration. While the consensus is that higher flexibility aids longevity, the trade-offs between material and dimensions of the probe and the best implantation technique remain uncertain. How does the damage caused by a silicon probe of minimal cross-section compare to that caused by a polyimide probe that is larger to harbor enough microelectrodes? Many of the forementioned studies addressed some of the sub-questions about device design in isolation, but, to our knowledge, no previous study directly compared the tissue compatibility of flexible and rigid electrode arrays of various widths and thickness that were either tethered or free floating. Knowledge about the relative influences of these factors on implant longevity will be essential for the future design choices for biocompatible brain implants.

We here present a comprehensive dataset of over a hundred brain slices collected from mice implanted with polyimide or silicon probes of various cross-sections, using either a tethered or untethered design. We developed a new quantitative histological analysis pipeline to investigate tissue reactions based on the presence of neurons, astrocytes, and microglia. Furthermore, we investigated the reaction along the entire probe, examining the cortical depth profile and underlying white matter. We observe a depth dependent immune response for all probes and report a general advantage of the flexible polyimide probes on tissue integrity, accompanied by weaker influences of other design parameters like probe dimensions.

## Results

We examined the influence of probe material, geometry, and implantation strategy on long-term tissue responses in the cortex of mice. We implanted rigid silicon probes and flexible polyimide probes that varied in shank width (35, 70, and 105 µm) and thickness (polyimide: 2, 5, 15 or 25 µm; silicon: 15, 25 or 50 µm), which were either tethered to the skull or left untethered. The probes had a comb-like design, consisting of three parallel shanks of identical thickness but different widths (Figure 1A-D). Many probes did not have functional electrodes because our main goal was to examine the tissue effects, but we also implanted probes with functional electrodes to assess recording quality over time. In total, 32 mice were implanted with three or four comb-like probes (i.e. a total of 9 to 12 shanks, Figure 1C), of which 30 mice were examined after approximately 6 months and two after 12 months of implantation. We inserted the probes with stereotaxic guidance, and we used temporary shuttle devices, which were retracted after insertion, to deliver the flexible probes. Tethered probes had wings that we attached to the skull with cement, and we placed untethered probes under a stainless-steel cap with a high inner roof that replaced a piece of the skull to prevent pressure from the skull onto the probes (Figure 1B). After the implantation period, we sacrificed and perfused the mice, sectioned their brains, and processed the sections for quantitative histology, recovering hundreds of probe shank tracks across the mice. We noticed that some of the shanks gave rise to loss of cortical tissue and quantified these lesions. Our histological analysis focused on neuronal density (NeuN), astrocytic reactivity (GFAP) and microglial activation (IBA1) (Figure 1C,D). In what follows, we will first quantify gross lesions caused by the probes in several mice. We will then characterize the immune responses across the cortical depth, the neuronal and glial reactions to the design parameters, the effects of implantation duration, and finally the relationship between tissue integration and the electrophysiological recording stability.

**FIGURE 1.**
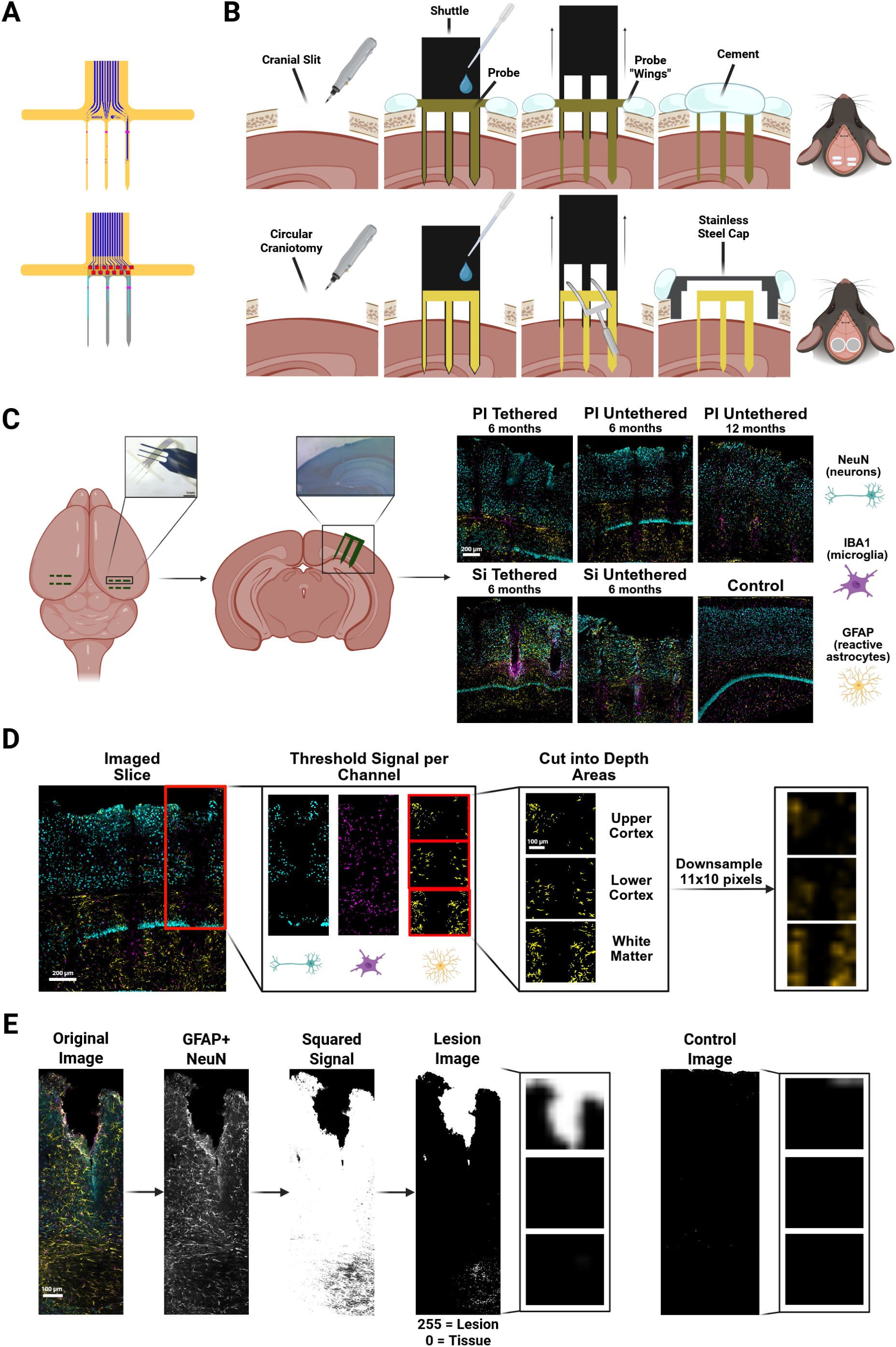
Implantation approach and quantification of the effects on the cortex. **(A)** Probe designs. Each probe consisted of 3 shanks of three different widths: 35, 70, and 105 µm and had one of several thicknesses. Polyimide probes (top) had a thickness of 2, 5, 15 or 25 µm and silicon probes (bottom) a thickness of 15, 25 or 50 µm. **(B)** Surgical procedures. We implanted 3-4 probes, with three shanks each in each mouse. Top: Tethered polyimide probes were inserted with a shuttle device after drilling a cranial slit of 1.5 mm into the mouse skull. The probes had wings that were attached to the skull with cement. The shuttle was removed and the craniotomy was sealed with cement. Bottom: Untethered polyimide probes were also inserted with a shuttle device. The probe detachment was supported with the help of a custom-made fork-like tool. A stainless-steel cap with a high inner roof was placed over the craniotomy and secured with cement. Silicon probe implantations followed the same procedure but did not require a shuttle device. **(C)** Immunohistochemistry. We sacrificed the mice after 6 or 12 months. We identified the lesions with Evan’s Blue (marking albumin) and identified neuronal density with NeuN, microglia with IBA1, and reactive astrocytes with GFAP. Right: Example images of tethered and untethered polyimide (PI) and silicon (Si) probes and a control image. Scale bar = 200 μm. **(D)** Quantification pipeline. We identified neurons, astrocytes, and microglia around each probe shank in three ROIs: Upper cortex, lower cortex and white matter. Scale bars represent either 200 or 100 μm. **(E)** We also identified regions with loss of cortical tissue and quantified the extent of the lesions. Right: example control image. Scale bar = 100 μm.

### Rigid probes cause more cortex loss

Several probes caused a loss of tissue, creating regions without cells (Figure 1E shows an example lesion). To quantify the tissue loss, we created binarized images of tissue vs. non-tissue (see Methods; Figure 1E). In short, we combined NeuN and GFAP fluorescence images and classified pixels as cortex or non-cortex. We then averaged the images per probe type and measured a normalized measure of tissue loss compared to unimplanted cortex, which we will call ΔTissueLoss. Higher values of ΔTissueLoss imply more tissue loss than control cortex (see Methods).

When we averaged across the entire depth profile, the tissue loss was more prominent for silicon probes than for polyimide probes (*z* = 4.21, *p* < .0001, *N_images_* = 400; Figure 2A). To account for differences in tissue reactivity and tissue stress along the probe trajectories, we defined three regions of interest (ROIs): (i) the upper cortex, (ii) the lower cortex and (iii) underlying white matter (Methods). Each ROI extended approximately 460 µm in depth and 400 µm in the horizontal direction. The upper cortical ROI started at the pial surface and included the superficial cortical layers, which are more directly affected by probe entry and disruption of the meninges than the lower cortex. The middle ROI comprised the deeper cortical layers. It was aligned on the white matter boundary and extended in the direction of the pia. The lowest ROI began at the cortical–white matter boundary and extended downward, only including white matter. These three ROIs are central to our quantification approach and will be used in the rest of the analysis.

**FIGURE 2.**
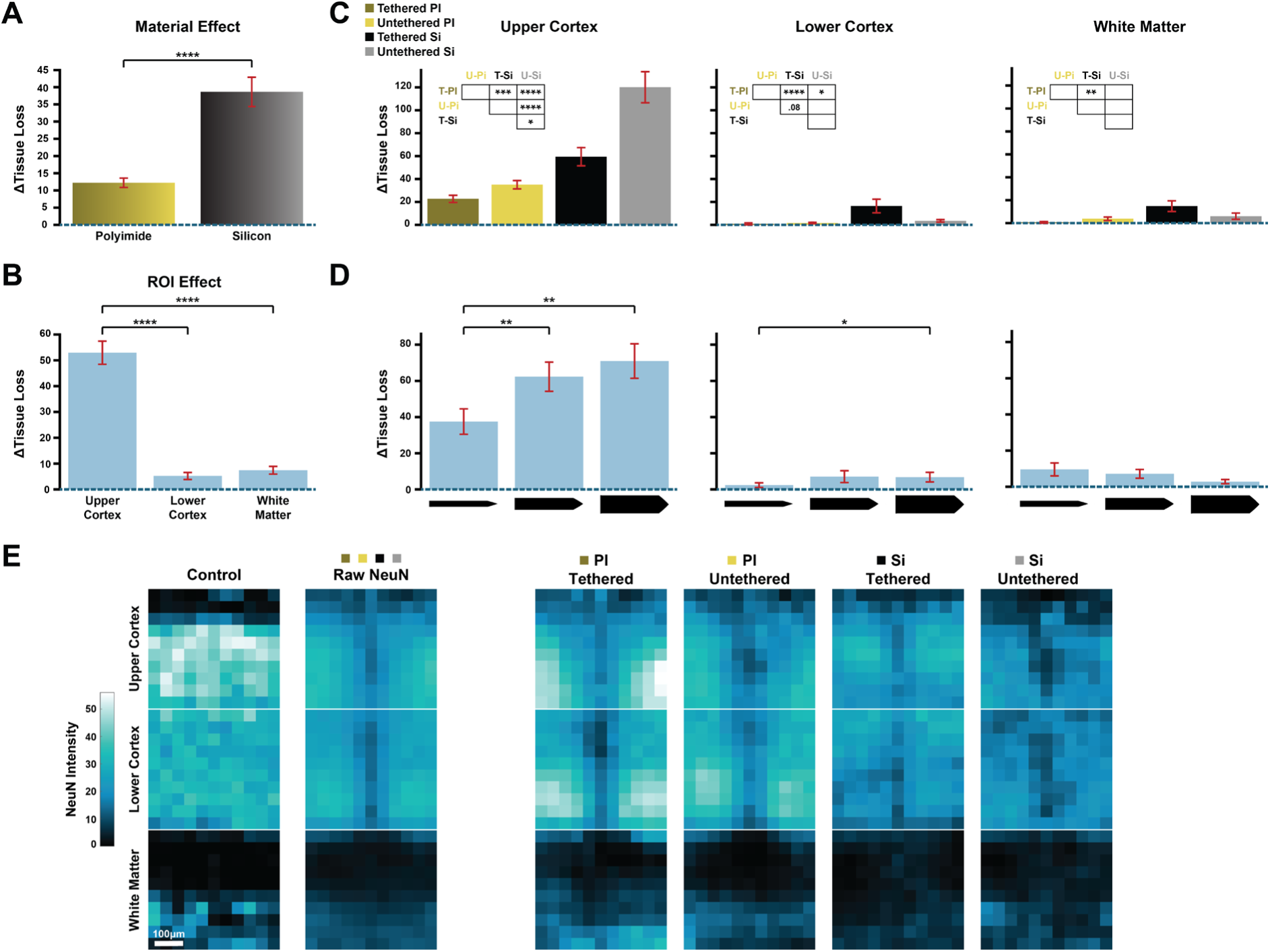
Loss of tissue caused by probe implantation, which was measured by comparing sections to those of control, non-implanted cortex. The signal ranges from 0 to a maximum of 255. Error bars represent SEM. * p < .05, ** p < .01, *** p < .001, **** p < .0001. **(A)** Material effect. Silicon probes caused more severe tissue loss than polyimide probes. **(B)** Region effect. Complete tissue loss was most pronounced in the upper cortex ROI. **(C)** Material and tethering effect in the three ROIs. T-PI = Tethered Polyimide; U-PI = Untethered Polyimide; T-Si = Tethered Silicon; U-Silicon = Untethered Silicon. **(D)** Effect of shank width in three ROIs. **(E)** NeuN signal after removing the pixels with complete loss of cortex. Left: control condition and average NeuN signal across conditions. Right: raw NeuN signal averaged over material/tethering conditions. NeuN binds to neurons, revealing neuronal loss in regions where the overall structure of the cortex was preserved. Scale bar = 100 μm.

Tissue loss was more pronounced in upper cortex than in the lower cortex (*z* = -12.8, *p*< .0001, *N* = 282) and the white matter (*z* = -11.9, *p* < .0001, *N* = 269; Figure 2B). We next examined the probe design parameters within the three ROIs (Figure 2C). In the upper cortex, untethered silicon probes caused more tissue loss than tethered polyimide (*z* = 6.88, *p* < .0001, *N* = 77), untethered polyimide (*z* = 4.56, *p* < .0001, *N* = 70) and tethered silicon probes (*z* = 2.98, *p* < .05, *N* = 72). Additionally, tethered silicon probes caused more tissue damage than tethered polyimide probes (*z* = 3.94, *p* < .001, *N* = 81). In the lower cortex, tethered polyimide probes caused the least amount of tissue loss, significantly less than tethered (*z* = 4.44, *p* < .0001, *N* = 61) and untethered silicon probes (*z* = 2.89, *p* < .05, *N* = 63). Tethered silicon probes also caused a loss of white matter, and more so than tethered polyimide probes (*z* = 3.61, *p* < .01, *N* = 51). To summarize, the probe material was a prime determinant of tissue loss, with silicon probes causing more damage than polyimide probes.

The width of the probe shanks also influenced the degree of tissue loss in the cortical ROIs (upper cortex: *χ*^2^(2) = 14.76, *p* < .001; lower cortex: *χ*^2^(2) = 6.53, *p* < .05). The 105 μm-wide shank caused more tissue loss than the 35 μm-wide shank in the upper (*z* = -3.45, *p* < .01, *N* = 98) and lower cortex (*z* = -2.55, *p* < .05, *N* = 80). Additionally, the 70 μm-wide shank caused more tissue loss than the 35 μm-wide shank in the upper cortex (*z* = -3.2, *p* < .01, *N* = 103; Figure 2D). We did not find significant effects of probe thickness on tissue loss.

We note, however, that tissue damage can also express as a more diffuse loss of neurons without a complete disruption of the cortex. We therefore quantified the density of remaining neurons with a NeuN stain. Figure 2E provides a first impression of how different probe types cause a decrease in neuronal density across cortical depth. Specifically, we converted the NeuN fluorescence images (Figure 1C,D) into quantitative profiles by creating a binary categorization of neurons in the image and down sampled the images to generate pixel wise representations along the probe track (see Methods). We then averaged the pixels across images in each condition. We will first describe the equivalent quantification for GFAP and IBA1 stains that measure the increases in astrocytic and microglia reactivity, before we return to the full quantification of the NeuN results, in a later section of the Results.

### Immune reactivity has a stereotypical depth profile

We will first examine the overall depth profile of the GFAP label by averaging across all the imaged shanks (*N* = 400 ROI images) and subtracting the label in non-implanted cortex. The ΔGFAP signal (Figure 3A) revealed a clear depth profile of astrocytic reactivity with two zones with a high signal. The first zone was in the superficial layers of the cortex, where the probes entered the tissue, and the second one at the boundary with the white matter (Figure 3A). We examined the average ΔGFAP profile within the upper cortical ROI by comparing the upper and lower half of this ROI. There was higher signal intensity at the tissue insertion site than in the bottom of the ROI (*t*(143) = 1.9, *p* < .05). A comparable analysis of the lower cortical ROI revealed a stronger ΔGFAP signal close to the white matter than in the upper region of this ROI (*t*(130) = -7.1, *p* < .0001). Although astrocytes are present in the white matter ROI of the control slices (middle panel in Figure 3A), probe insertions caused increased GFAP signal at the border with the cortex, which was stronger than in the lower part of the white matter ROI (*t*(117) = 3.3, *p* < .0001).

**FIGURE 3.**
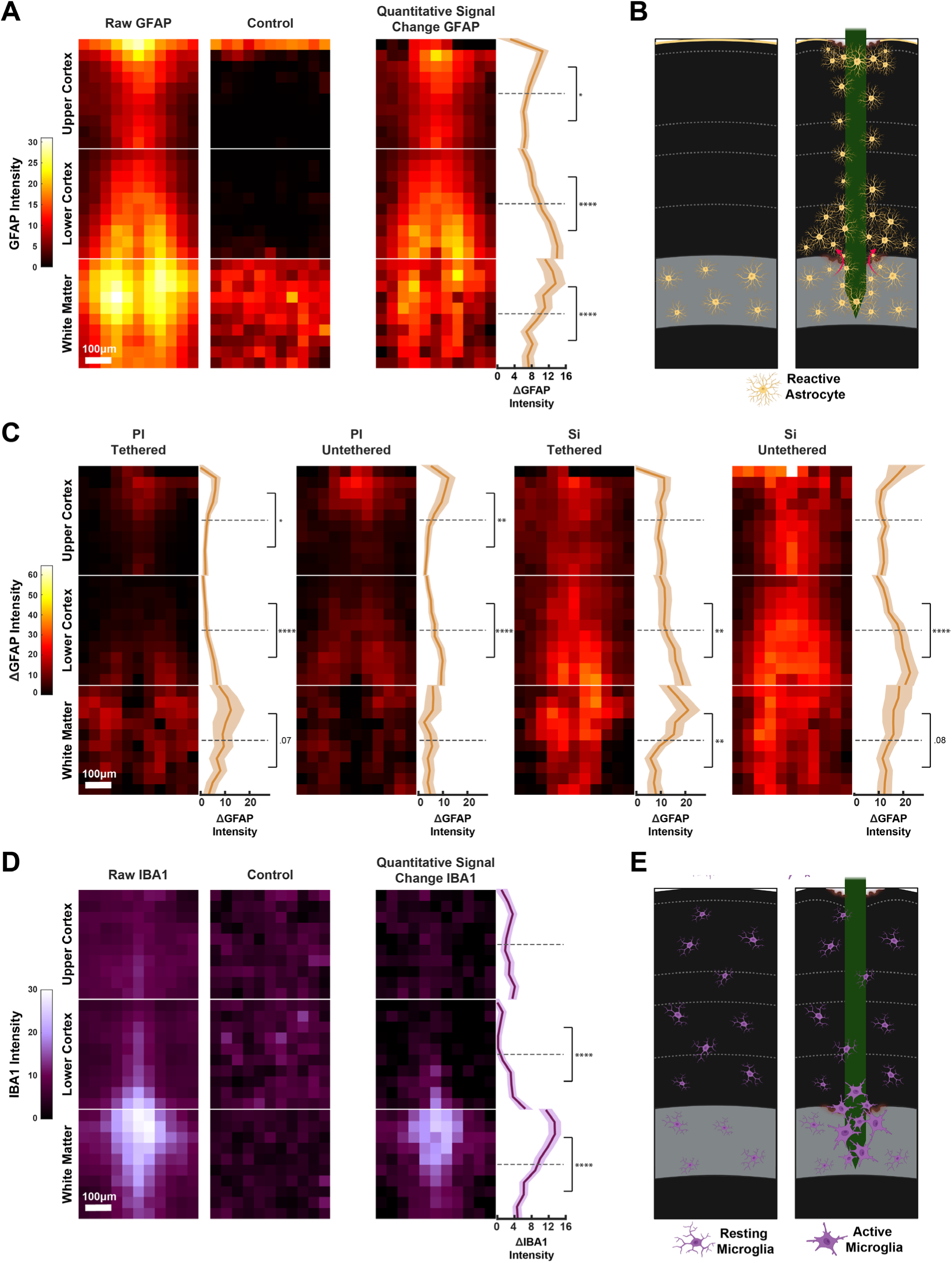
Depth profile of the immune responses. Scale bar = 100 μm. **(A)** Average astrocyte response across all experimental conditions. Left: GFAP signal across all probes. Middle: Control GFAP signal. Right: Difference (ΔGFAP). Line graphs display the mean ΔGFAP along cortical depth (shaded area SEM). We compared ΔGFAP between the upper and lower half of each ROI. * p < .05, ** p < .01, *** p < .001, **** p < .0001. **(B)** Illustration of the astrocytic response across cortical depth. Left: Untreated tissue. Right: Implanted tissue. **(C)** Astrocytic depth responses for individual material and tether conditions. PI = polyimide, Si = Silicon. There is an increase in GFAP at the boundary between cortex and white matter for every probe type. **(D)** Average microglia response (quantified with IBA1) for probes (left), control sections (middle) and the difference (right). Differences within region parts show an increase of microglia in the lower cortex and white matter at their intersection. **(E)** Illustration of microglia reaction across depth. Left: Untreated tissue. Right: Implanted tissue.

We next examined the influence of electrode material and tethering, pooling across probe dimensions in each of the conditions. Polyimide probes caused a higher GFAP signal at the tissue insertion site (Polyimide (PI) Tethered: *t*(42) = 2, *p* < .05; PI Untethered: *t*(35) = 3.3, *p* < .01) and close to the white matter (PI Tethered: *t*(29) = -4.58, *p* < .0001; PI Untethered: *t*(36) = -5.18, *p* < .0001). In contrast, silicon probes caused a more general GFAP signal elevation throughout cortical depth, while still displaying a higher signal increase at the boundary to the white matter (Silicon (Si) Tethered: *t*(30) = -2.59, *p* < .01; Si Untethered: *t*(32) = -4.25, *p* < .0001) and, unlike for polyimide, also in the white matter itself (*t*(30) = 3.1, *p* < .01) (Figure 3C).

To assess whether these astrocytic responses were accompanied by the activation of microglia, we next examined IBA1 staining. IBA1 is a marker that is expressed by both resting and activated microglia that is used to quantify microglial recruitment following neural injury and implantation (Potter et al., 2012; Szarowski et al., 2003). When we averaged across all the shanks (*N* = 332 ROI images), we observed increase in IBA1 signal intensity at the border between the cortex and the white matter (Figure 3D,E). When we divided the lower cortical ROI into two equal compartments based on depth, the IBA1 signal was stronger in the lower than in the upper half (*t*(116) = -6.2, *p* < .0001). Within white matter, the increased microglia reaction occurred just below the cortex (*t*(99) = 6.3, *p* < .0001). We did not observe large IBA1 differences between shanks of silicon or polyimide, and also only weak effects of tethering (Suppl. Figure 3).

### Polyimide probes display better tissue integration than silicon probes

In the above, we described the pixel-by-pixel depth profiles of NeuN, GFAP and IBA1. We will now turn to the main goal of our study: a systematic analysis of the effect of the width, thickness and tethering of flexible and stiff probes on ΔNeuN, ΔGFAP, and ΔIBA1. For this we concentrate our analyses on average signal values per ROI (see Methods).

We applied ANOVAs for the three markers and will describe the significant main effects (see Methods for details). The first main effect reflected differences in expression between the three ROIs, for each of the markers ΔNeuN (*F*(1,203) = 36.1, *p* < .0001), ΔGFAP (*F*(2,290) = 11.66, *p* < .0001), and ΔIBA1 (*F*(2,227) = 40.49, *p* < .0001). ΔNeuN was larger in the upper cortex (post-hoc t-test, *t*(277) = -4.9, *p* < .0001) than in the lower cortex (Figure 4A top). In contrast, ΔGFAP was larger in the lower cortex than in the upper cortex (*z* = 5.05, *p* < .0001, *N* = 282) and white matter (*z* = -2.59, *p* < .05, *N* = 269) (Figure 4A middle). ΔIBA1 in the white matter was larger than in the cortical ROIs (upper cortex vs. white matter: *z* = 7.95, *p* < .0001, *N* = 215; lower cortex vs. WM: *z* = 7.42, *p* < .0001, *N* = 217) (Figure 4A bottom).

**FIGURE 4.**
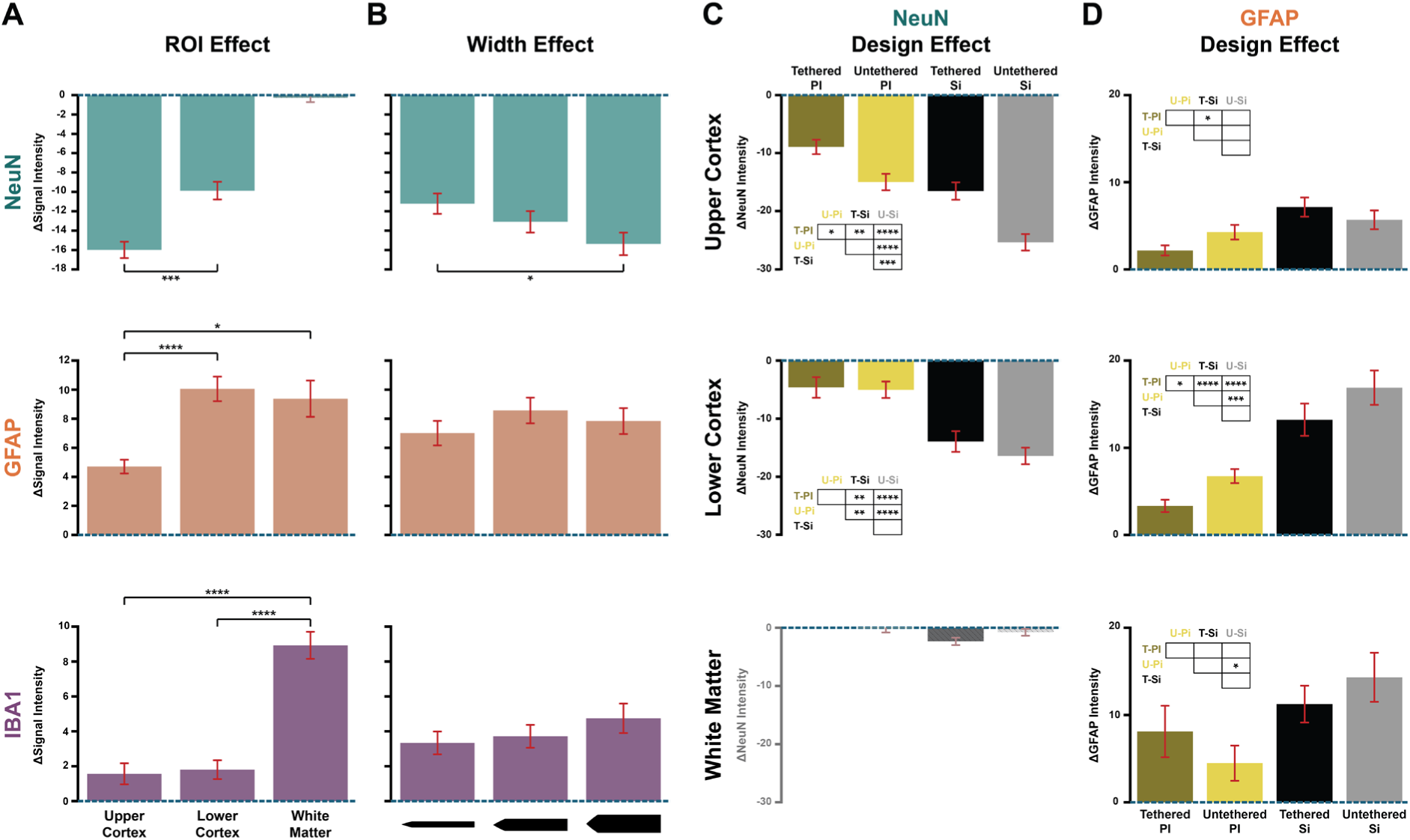
Quantitative analysis of NeuN, GFAP and IBA1 signals. Neuronal loss, astrocyte and microglia response are calculated as ΔNeuN, ΔGFAP, and ΔIBA1 by taking the difference with the control condition. **(A)** ROI effects. Top: Neuronal loss quantified as ΔNeuN. Middle: Astrocytic response. Bottom: Microglia response. * p < .05, ** p < .01, *** p < .001, **** p < .0001. **(B)** Width effect. Top: Neuronal loss for the three shank widths (35, 70, 105 µm) across ROIs. Middle: Astrocytic response. Bottom: Microglia response. **(C)** Probe design effect on neuronal loss in each ROI. Top: Neuronal loss caused by material and tether choices in the upper cortex. Middle: Lower cortex. Bottom: White matter. Given the low number of neurons, we did not conduct a statistical analysis for the white matter data. T-PI = Tethered Polyimide; U-PI = Untethered Polyimide; T-Si = Tethered Silicon; U-Silicon = Untethered Silicon. **(D)** Probe design effect on astrocytic response.

We obtained a main effect of shank width on ΔNeuN (*F*(2, 203) = 10.11, *p* < .01), with higher neuronal loss around the 105 μm-wide shank than the 35 μm-wide shank (*t*(170) = -2.64, *p* < .05) (Figure 4B, top). Shank width had no main effect on the intensity of the immune response (ΔGFAP and ΔIBA1; Figure 4B). We found an interaction between probe material and shank width in the ΔIBA1 signal (*F*(2,227) = 3.59, *p* < .05), which was caused by an effect of the shank width of silicon probes on ΔIBA1 (105 μm-wide vs. 35 μm *z* = -2.45, *p* < .05, *N* = 81).

We will now describe the effects of tethering and material (Figure 4C,D). There was a main effect of material on the ΔNeuN (*F*(1,203) = 80.16, *p* < .0001) and ΔGFAP (*F*(1,290) = 48.44, *p* < .0001), with more neuronal loss (*t*(277) = 8.32, *p* < .0001) and higher astrocytic reactions (*z* = 7.12, *p* < .0001, *N* = 400) for silicon probes. There also was a main effect of tethering on ΔNeuN (*F*(1,203) = 14.72, *p* < .001), because untethered probes caused more neuronal loss than tethered probes (*t*(277) = 3.28, *p* < .01). Tethering did not have a significant effect on the immune response as quantified by ΔGFAP and ΔIBA1.

We further examined material-tethering interactions within each ROI, first focusing on the loss of neurons with ΔNeuN. In the upper cortex, the tethered polyimide probes caused less neuronal loss than the untethered polyimide probes (*t*(78) = 3.19, *p* < .05), the tethered silicon probes (*t*(78) = 3.87, *p* < .001), and the untethered silicon probes (*t*(76) = 8.64, *p* < .0001) (Figure 4C, top). Untethered silicon probes caused more neuronal loss than untethered polyimide (*t*(69) = 5.16, *p* < .0001) and tethered silicon probes (*t*(69) = 4.25, *p* < .001). In the lower cortex, the tethering effect of both polyimide and silicon probes was not present, but polyimide probes caused less neuronal loss than the silicon probes (*t*(128) = 6.55, *p* < .0001) (Figure 4C, middle).

We carried out the equivalent analysis for ΔGFAP to measure immune responses (Figure 4D). In the upper cortex, the tethered silicon probes caused a higher astrocytic reaction than the tethered polyimide probes (*z* = 3.92, *p* < .001, *N* = 81) (Figure 4D top). In the lower cortical ROI there were many differences between conditions, which are summarized in Figure 4D middle. The tethered polyimide probes caused the least amount of astrocytic reaction, with significantly lower ΔGFAP than the untethered polyimide (*z* = 2.77, *p* < .05, *N* = 67), tethered silicon (*z* = 5.1, *p* < .0001, *N* = 61) and untethered silicon probes (*z* = 6.57, *p* < .0001, *N* = 63). Furthermore, untethered polyimide probes caused a lower ΔGFAP than untethered silicon probes (*z* = 4.08, *p* < .001, *N* = 70). The tethering effect was not significant for the silicon probes. In the white matter, untethered polyimide probes caused the smallest ΔGFAP and untethered silicon probes the largest ΔGFAP, a difference that was significant (*z* = 3, *p* < .05, *N* = 67) (Figure 4D bottom). We did not find significant effects of material or tethering on ΔIBA1 in the three ROIs.

As a measure for the overall tissue reaction, we defined Idx_TissueReaction_, based on ΔNeuN and ΔGFAP (see Methods).

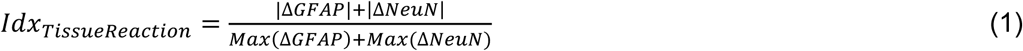

This index combines the astrocytic reaction with the neuronal loss. Because ΔNeuN tends to be negative and ΔGFAP positive we took the absolute values and normalized to the sum of maximal values (range from 0 to 255) so that Idx_TissueReaction_ ranges from 0 (no tissue reaction) to 1 (maximal astrocytic response and loss of neuronal signal).

The GFAP and NeuN results described so far predict a higher index for silicon probes, and this is what we observed. Idx_TissueReaction_ was larger for silicon than for polyimide in both the upper (*z* = 6.41, *p* < .0001, *N* = 149) and lower cortex (*z* = 7.75, *p* < .0001, *N* = 130). Unexpectedly, untethered probes caused a higher Idx_TissueReaction_ in the upper cortex than tethered probes (*z* = 4.26, *p* < .0001, *N* = 149). The width of the shanks also influenced Idx_TissueReaction_ (*χ*^2^(2) = 11.21, *p* < .01). Pairwise tests revealed significant differences between the 35 μm and 105 μm shanks in the lower cortex for silicon probes (*z* = -2.29, *p* < .05, *N* = 38). Figure 5A presents an overview of Idx_TissueReaction_ across design variables. In this figure color represents material and tethering, the size of the symbols probe thickness and there are separate rows for probe width and ROI.

**FIGURE 5.**
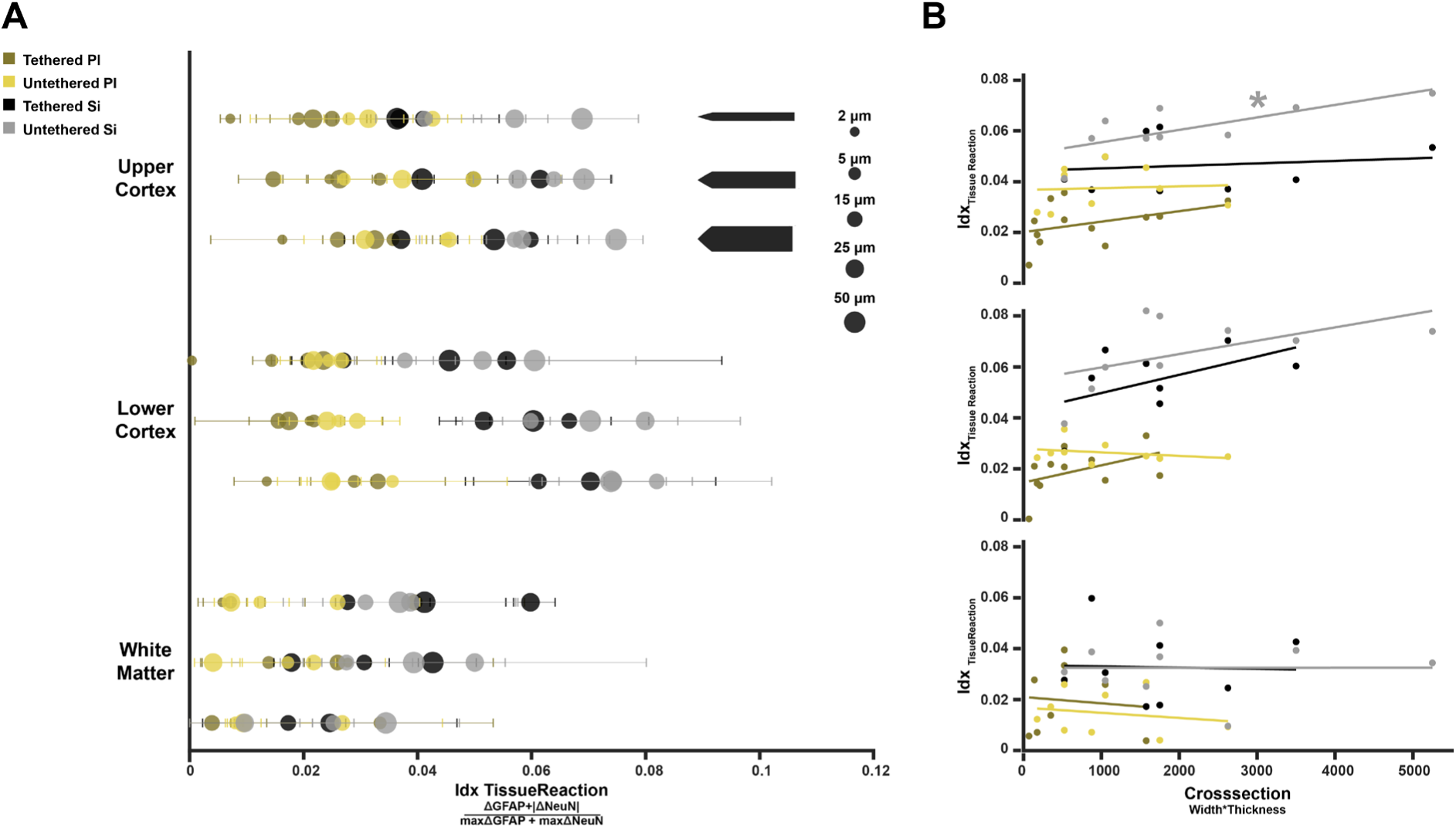
Idx_TissueReaction_ is the combined signal of ΔNeuN and ΔGFAP. **(A)** Idx_TissueReaction_ for all experimental conditions per ROI. Top: Upper cortex. Colors indicate material and tethering and circle sizes probe thicknesses. Rows denote shank widths (from top to bottom: 35, 70, 105 µm). Middle: Lower cortex. Bottom: White Matter. No statistical analysis was performed on Idx_TissueReaction_ in white matter, because NeuN is non-informative so that the index mainly reflects GFAP. **(B)** Effect of probe cross-section on tissue reaction. Top: Upper cortex, Middle: Lower cortex, Bottom: White matter. * p < .05, ** p < .01, *** p < .001, **** p < .0001.

### The relation between probe cross-section and the tissue reaction

We investigated the full effect of probe dimension on Idx_TissueReaction_ based on the cross-section of the shanks (width x thickness). We focused our data on the cortical ROIs, excluding white matter, and fitted a linear regression model with material, tethering, ROI and cross-section as predictors. The model explained variation in Idx_TissueReaction_ (*F*(279,274) = 48.7, p < .0001). Consistent with the previous results, material and tethering increased Idx_TissueReaction_, with more tissue damage for silicon (*ꞵ* = .02, *t*(274) = 10.03, p < .0001) and untethered probes (*ꞵ* = -.01, *t*(274) = -4.46, p < .0001). Furthermore, larger cross-sections increased Idx_TissueReaction_ (*ꞵ* = 3.1·10^-6^, *t*(274) = 2.7, p < .01). We next used separate linear regressions to test the influence of cross-section on average Idx_TissueReaction_ for each material-tethering probe combination and ROI (Figure 5B). In this analysis, the only significant influence of cross-section occurred for the untethered silicon probes in the upper cortex (*t*(7) = 2.95, *p* < .05). Hence the analysis of Idx_TissueReaction_ replicated the effects of material and tethering, and it also revealed a significant, yet smaller influence of shank cross-section on tissue damage.

### Prolonged implantation is associated with additional loss of neurons

We obtained the results described so far with an implantation duration of 6 months. We additionally implanted 15 μm and 25 μm-thick untethered polyimide probes for 12 months to examine the effect of a longer implantation duration (Figure 6A). We found fewer cases with complete tissue loss in the upper cortex for the 12-month than in the 6-month group (*z* = -2.11, *p* < .05, *N* = 45) (Figure 6B). However, we observed a larger ΔNeuN in the lower cortex for probes for the longer implantation duration, indicative of loss of neurons (*t*(45) = 4.94, *p* < .001) (Figure 6C, left). We did not find significant effects of implantation duration on the immune response as measured with ΔGFAP and ΔIBA1 (Figure 6C, middle and right). When we assessed Idx_TissueReaction_, we found that tissue reaction was larger in the 12-month than in the 6-month group (*z* = 3.24, *p* < .01, *N* = 92). Hence, the longer implantation duration gave rise to a slightly larger loss of neurons in the lower cortical layers, which was presumably also reflected by an increase in Idx_TissueReaction_.

**FIGURE 6.**
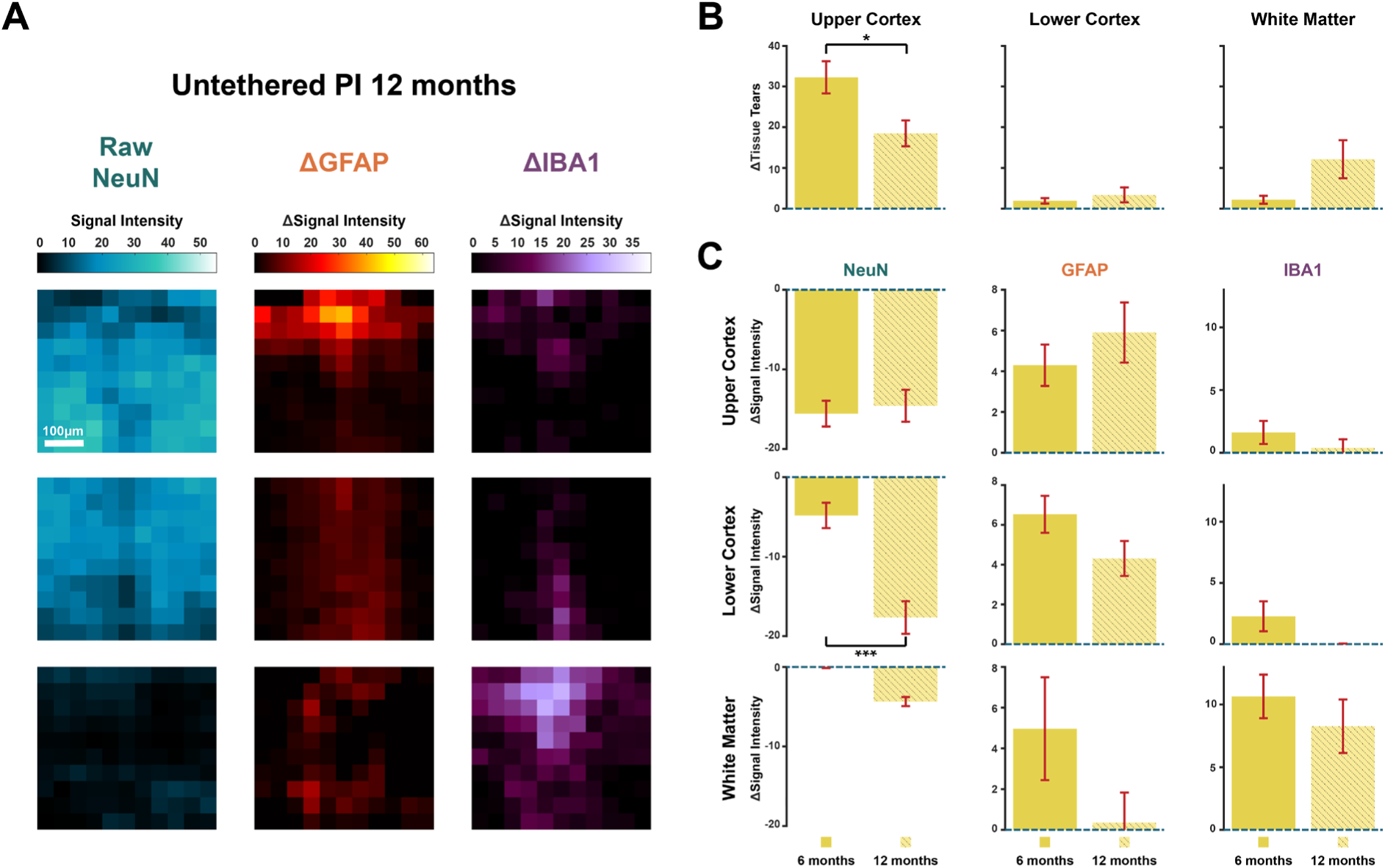
Implantation duration effect of untethered polyimide probes. **(A)** Average neuronal, astrocytic, and microglial signals for probes implanted for 12 months. Left to right: Raw NeuN signal intensity displaying neurons, ΔGFAP marking astrocytic activity and ΔIBA1 signal marking microglial activity. Top to bottom: Upper cortex, lower cortex, and white matter. Scale bar = 100 μm. **(B)** Tissue loss caused by untethered polyimide probes implanted for 6 or 12 months. Left to right: Upper cortex, lower cortex, and white matter. **(C)** Comparison of ΔNeuN (left), ΔGFAP (middle column) and ΔIBA1 (right) in upper cortex (top), lower cortex (middle row), and white matter (bottom). Error-bars depict SEM. * p < .05, *** p < .001.

### Polyimide probes exhibit better longitudinal recording quality

To test the effects of probe design choices on the electrophysiological recording quality, we analyzed the neuronal data of 2 mice implanted with polyimide probes with thicknesses of 5, 15, and 25 μm, as well as 3 mice implanted with silicon probes with thicknesses of 15, 25, and 50 μm. The probes were implanted in the left and right primary visual cortex, and we recorded neuronal signals while mice passively viewed a checkerboard stimulus (see Methods, Figure 7A). We collected electrophysiological data for 24 weeks post implantation and calculated the quality of the visual response as signal-to-noise ratio (SNR) for each electrode (*N*_PI_ = 42, *N*_Si_ = 81). The SNR was defined as the average peak neuronal response to the checkerboard, divided by the standard deviation of pre-stimulus baseline activity across trials (see Methods). We observed a decrease in the quality of the signal over time for both polyimide and silicon probes (PI: *t*(1678) = -12.3, *p* < .0001; Si: *t*(3238) = -22.3, *p* < .0001) (Figure 7B). To investigate design effects on the recording quality, we sorted the data into 6 time-bins of one month. From the first month onward, polyimide probes displayed a better recording quality than silicon probes (*z* = -3.14, *p* < .01, *N* = 123). This difference persisted until the sixth month (Figure 7C). We did not find consistent effects of probe thickness or shank width on the recording quality (Suppl. Figure 5A). There was a slightly better signal for the 50 µm than for the 15 μm-thick silicon probes in the first month (*z* = 2.61, *p* < .05, *N* = 54) as well as a better signal for 5 μm than 25 μm-thick polyimide probes in the fifth month (*z* = -2.45, *p* < .05, *N* = 33) (Suppl. Figure 5A).

**FIGURE 7.**
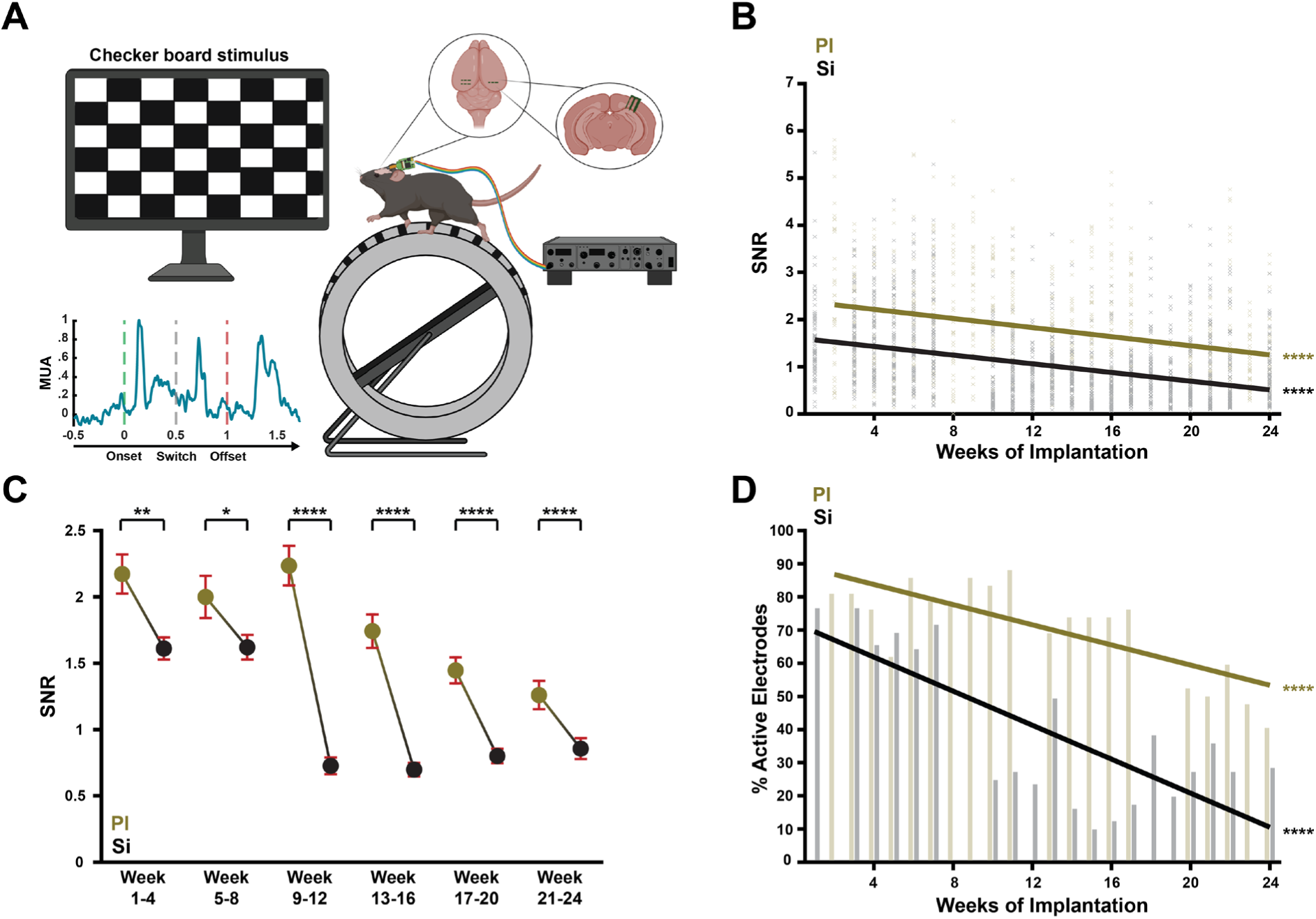
Electrophysiological recordings of V1 activity elicited by visual stimuli, recorded with polyimide and silicon probes. **(A)** Experimental set-up. We implanted with 3 polyimide or silicon probes in the primary visual cortex of mice. The mice saw a checker-board stimulus that reversed after 500ms and disappeared after 1 s. We computed the signal-to-noise ratio (SNR) as measure of the crispness of the visual response in the multiunit activity. **(B)** SNR of all electrodes on polyimide and silicon shanks across 24 weeks of implantation. Both materials displayed a loss of recording quality over time. **(C)** Comparison of polyimide and silicon electrode SNRs in six time-bins. Across the entire experimental duration, electrodes on PI probes displayed a better recording quality than those on Si probes. **(D)** Percentage of useful electrodes (SNR > 1) on PI and Si probes over time. * p < .05, ** p < .01, *** p < .001, **** p < .0001.

We next examined the percentage of electrodes with useful signal and chose an arbitrary cut-off of SNR > 1. In the first recording week, we started with 82% useful electrodes on polyimide probes and 77% on silicon probes. In the last recording week, these percentages had decreased to 40% for polyimide and 28% for silicon probes. Hence, the percentage of useful electrodes decreased for both materials, but the decrease in useful electrodes was steeper for silicon probes (Si: *ꞵ* = -2.57, *t*(18) = -4.98, *p* < .0001; PI: *ꞵ* = -1.52, *t*(18) = -4.99, *p* < .0001). We note that the drop in useful electrodes in polyimide probes could have been caused by an infection at the implantation site in one of the two mice.

## Discussion

We present a new quantitative histological analysis pipeline to systematically measure the influence of material and design choices of implantable intracortical probes on brain tissue. We investigated three major adverse effects on brain tissue: loss of neurons, glial scarring and inflammatory responses. We systematically varied several probe dimensions and used a rigid and a flexible shank material, while systematically and quantitatively examining hundreds of brain slices. Thereby the present study is, to our knowledge, completer and more extensive than previous foundational studies in this domain.

The central finding is the superior tissue compatibility of polyimide probes compared to silicon probes. Polyimide probes caused fewer cortical lesions, preserved a higher density of neurons around the implanted shanks and elicited a weaker astrocytic reaction than silicon probes. Furthermore, the longer implantation of 12 months only weakly increased the neuronal loss over that in the 6 months group. Limiting glial scarring while maintaining close proximity between electrodes and healthy neurons is critical for the long-term functionality of neural interfaces (Otte et al., 2022; Polikov et al., 2005; Szarowski et al., 2003), as a shorter electrode-neuron distance improves recording fidelity and stimulation efficacy (Buzsáki, 2004). Consistent with these principles, polyimide probes maintained a better recording quality than silicon probes. Furthermore, the late-stage reduction of the fraction of useful electrodes on polyimide probes was presumably attributable to an infection at the implantation site in one of the mice, which led to extensive gliosis that encapsulated all implanted devices. The complication likely arose from surgical factors rather than from the polyimide material itself, as prior studies demonstrated exceptional long-term stability of polyimide probes (Luan et al., 2017; Zhao et al., 2023) and we did not observe a higher incidence of infections for polyimide probes than silicon probes across the full cohort.

Larger shank cross-sections increased the tissue damage in the cortex, although the effect was smaller than those for material and tethering (Figure 5B). Probe cross-section is thought to determine the extent of neural and vascular disruption during implantation (Seymour & Kipke, 2007; Stice et al., 2007), although larger probes do not always impair recording quality (Vomero et al., 2022). In our data, probe thickness alone neither influenced neuronal loss nor the immune response for either material. In contrast, increasing shank width (35 vs. 105 μm) caused more neuronal loss, evident as larger cortical lesions and reduced neuronal density, but it did not amplify astrocytic or microglial activation. When comparing the effects of shank width and thickness, it is important to consider the mechanical coupling, which is asymmetric across shank dimensions. The three parallel shanks were connected along the width dimension but not along the thickness dimension, which may have increased stiffness laterally and thereby have contributed to the stronger effect of shank width. Our probes had an inter-shank spacing of 400 µm and it is conceivable that different spacings may change the balance between the influence of thickness and width.

A key novelty of our study is the investigation of tissue reactions along the entire depth profile of the chronically implanted devices. Examining astrocytic reactivity along the probe trajectory revealed two prominent foci of reactive gliosis: at cortical surface where the probe enters the tissue and at the boundary between the cortex and the white matter. The elevated response at the cortical surface is likely caused by disruption of the meninges and the displacement of fibroblasts into the cortex. Previous work has shown that persistent contact between neural probes and ruptured meninges induces a sustained glial and fibroblastic reaction (Kim et al., 2004; Woolley et al., 2013). In principle, this reaction could be reduced by fully embedding the probe within the brain tissue, as demonstrated for single shank designs by Kim et al. (2004). However, such an approach is impractical for multi-shank probes, where complete immersion of the connecting backbone would likely exacerbate tissue damage.

We also observed a pronounced astrocytic response at the cortex-white matter boundary. Because astrocytes are naturally abundant in white matter, penetration into this region may also cause an astrocytic “spillover” effect, elevating astrocytic reactivity in the adjacent lower cortical layers. Importantly, this response was superimposed on a more general astrocytic reaction to the probes, as the GFAP level in the white matter itself also increased around the shanks. Hence, the astrocytic response along the probe exhibited a U-shaped profile within the cortex, with a comparatively low number of reactive astrocytes in the middle cortical layers, which are usually targeted for electrophysiological recording and stimulation.

Microglial responses exhibited a distinct depth dependence, with a pronounced IBA1 elevation at the boundary between cortex and white matter that persisted even after 12 months of implantation. Although microglia are primarily associated with the acute immune response to probe implantation (Kozai et al., 2012; Potter et al., 2012), they can remain chronically activated around the implant because the probes are not cleared by phagocytosis (Otte et al., 2022; Potter et al., 2012). Importantly, microglia may not be inherently detrimental, as they can adopt both pro- and anti-inflammatory phenotypes (Cherry et al., 2014) and their presence may not impair recording performance (Prasad et al., 2014). Because white matter is usually not targeted for recording and stimulation, the activation of microglia in the white matter will not impair device functionality. However, it is uncertain whether the microglial response in white matter could exert longer-term influences on the nearby cortical tissue. Hence, it may be beneficial to avoid penetration into the white matter if the goal is to interface with the cortex. We compared two implantation strategies: a tethered configuration, in which probes were fixed to the skull, and an untethered figuration, in which probes were intended to reside more freely within the cortex. In contrast to previous studies suggesting improved tissue integration for untethered devices (Kim et al., 2004; Vomero et al., 2022), we observed a higher tissue reaction for untethered probes, particularly in the upper cortex. We hypothesize that this discrepancy may reflect a difference in surgical approach rather than an inherent disadvantage for untethered designs. Specifically, our untethered devices were associated with a larger surgical footprint, because we used a 3-mm-diameter craniotomy rather than the narrow cranial slit of the tethered devices. In several cases, edema occurred in the craniotomy of untethered probes, potentially resulting from the high inner roof of the protective cap used to close the skull. This configuration may have allowed mechanical interactions between the probes and the cap roof (Suppl. Figure 6). Nevertheless, during brain explantation most untethered probes remained fully embedded in the cortex, arguing against inadvertent tethering to the cap.

Overall, our findings highlight the importance of electrode material for long-term tissue integration. Within the range of probe designs examined here, flexible probes consistently showed advantages over rigid ones, whereas the effects of probe dimensions were weaker. This observation nuances the idea that slim but rigid devices are necessarily better accepted by the tissue than larger yet flexible probes (Otte et al., 2022). Importantly, our findings suggest that further miniaturization is valuable, but that the advantages need to be balanced with the functional requirements.

From a practical perspective, moderate increases in shank width or thickness may offer design advantages at the cost of only a moderate increase in tissue damage. For example, a shank with a width of 70 μm allows more tracks and electrodes than one of 35 μm without causing much more tissue damage. Similarly, although ultra-thin probes demonstrated excellent tissue integration and recording qualities (Luan et al., 2017; Zhao et al., 2023), the effect of probe thickness did not reach statistical significance here. Hence, probes with a thickness of ∼10 μm may represent a practical and biologically acceptable compromise, offering improved manufacturability, mechanical robustness, reduced cross-talk (Porto Cruz et al., 2025), and a higher implantation success rate than ultra-thin devices.

Finally, despite their less favorable tissue response, rigid probes retain important advantages, particularly for the integration CMOS-based functionality. Our results suggest that slightly bulkier devices might achieve tissue compatibility that is almost as good as ultra-thin probes, provided they remain flexible. This finding could inspire hybrid design strategies, by for example embedding rigid CMOS components within flexible substrates, as a promising path forward (De Dorigo et al., 2018) for combining advanced functionality with a favorable long-term tissue compatibility.

## Methods

### Animals

We implanted a total 32 C57BL/6J mice (12 female, 20 male) at 12-14 weeks of age. Of these, 10 mice (5 female, 5 male) received 4 non-functional tethered or untethered flexible polyimide probe combs for 6 months. Additionally, 10 mice (5 female, 5 male) received 3 non-functional tethered or untethered rigid silicon probe combs for 6 months. We implanted three untethered flexible polyimide arrays for 12 months in two male mice. These mice were housed socially in a normal daylight cycle. In the remaining 10 mice (2 female, 8 male) we implanted functional polyimide or silicon electrode arrays for electrophysiology. Due to animal health issues or complications with the connector, we did not obtain useful data in three mice with functional polyimide probes and two mice with functional silicon probes. The mice in these groups were housed in a reversed daylight cycle and solitary to protect the integrity of the implant. The study protocol (AVD-801002016631) was approved by the CCD (Central Commissie Dierproeven) and the ethical committee of the Royal Netherlands Academy of Arts and Sciences.

### Design of the probes

We used flexible polyimide-based and rigid silicon-based probes with the shape of a small comb (Suppl. Figure 1). All probes consisted of three shanks with a width of 35, 70 and 105 µm and an inter-shank spacing of 400 µm. We used four thicknesses, 2, 5, 15, and 25 μm, for the polyimide probes and three thicknesses, 15, 25, and 50 μm, for the silicon probes.

We used several types of probes. The first type were untethered probes (Suppl. Figure 1A-ii and C) made of polyimide or silicon. The second type were tethered because the probe comb of polyimide (Suppl. Figure 1A-iii) or silicon (Suppl. Figure 1B-ii and Suppl. Figure 1D) was connected to a 5-mm-long cable of polyimide. The third type were functional polyimide (Suppl. Figure 1 A-i) and silicon probes (Suppl. Figure 1C) with integrated micro electrodes which could be connected via a polyimide cable with 30 µm wide metal tracks (Suppl. Figure 1B-I and Suppl. Figure 1D) to a ZIF interface that could be attached via a ZIF-CLIP connector to a Tucker Davis Technologies headstage during recordings. The electrodes were 15 × 15 µm^2^ in size and they had a linear arrangement with a pitch of 40 µm and the tracks in the shanks had a width of 3 µm. The electrodes were coated with iridium oxide (IrO_x_) to decrease the electrical impedance.

Close to the probe base, we included visible markers with a length of 50 µm and width which was 13 µm smaller than the shank to be able to determine the appropriate implantation depth during surgery. The tethered probes had wings of 250 µm × 1500 µm that were fixed to the skull and were absent from the untethered probes. Most probes were non-functional and only consisted of the part in the brain without electrical wiring. For probe implantation, we mounted the flexible polyimide probes onto silicon shuttles (Suppl. Figure 1E) and used bio-dissolvable polyethylene glycol (PEG) as adhesive to facilitate insertion. The PEG dissolved when it came in contact with moisture in the brain, allowing the retraction of the shuttle device after probe implantation (Figure 1B).

### Probe manufacturing

The fabrication processes of the probe test structures and functional neural probes based on polyimide (PI) and silicon (Si) are summarized in Suppl. Figure 2. As shown in Suppl. Figure 2A, the PI-based neural probes are realized on Si substrates. The process applies spin-coating and imidization of a first PI layer, and deposition and patterning of the metallization layer interfacing electrode sites and contact pads using sputter deposition of platinum and a lift-off process (Böhler et al., 2023; Lewis et al., 2024). Individual polyimide layer thicknesses between 2.5 and 5 µm were adjusted by the spin-coating parameters spin speed and duration, whereas thicker polyimide layers (i.e. 12.5 µm) are realized by repeated coating/imidization cycles. Platinum deposition is followed by spin-coating a second polyimide layer which is patterned by reactive ion etching (RIE) in an oxygen plasma to define the electrode sites with a small via with a diameter of 8 µm and the contact pads of the ZIF interface. Next, the 15×15 µm^2^electrodes were defined by sputter deposition and lift-off patterning of IrO_x_. Finally, the geometrical shape of the polyimide based neural probes with common base and three shanks of different width is defined by RIE. The probes were peeled manually from the Si substrate using tweezers. The polyimide based untethered and tethered test structures were fabricated similarly. We spin-coated up to 5 polyimide layers, depending on the requested polyimide layer thickness.

The silicon-based neural probes are manufactured using established processes, as described in previous studies (Barz et al., 2017; Guardamagna et al., 2022; Herwik et al., 2009; Michon et al., 2016; Seidl et al., 2011) and the processing steps have been summarized in Suppl. Figure 2C. First, an insulation layer of silicon oxide (SiO_x_) and silicon nitride (Si_x_N_y_) was deposited using plasma enhanced chemical vapor deposition (PECVD). The metallization of pads, interconnects and recording sites was deposited and patterned using sputtering and a lift-off process. The metallization is insulated by a passivation layer, again deposited by PECVD. Next, the insulation and passivation stack was patterned by RIE to expose the contact pads on the probe base. At the position of the recording sites vias were defined to access individual metal tracks. Like for the polyimide-based neural probes, the electrodes consist of IrO_x_ which is patterned by lift-off. The probe shape is realized by a combination of RIE and deep RIE (DRIE) using the Bosch process (Laermer & Urban, 2003). The probes were thinned and released using the so-called etching-before-and-after-grinding (EBAG) process. The process has been introduced by Sharma et al. (2023) and is based on the etching-before-grinding (EBG) process, initially proposed by Herwik et al. (2011). After wafer grinding to an initial thickness of 70 µm, the probe shanks are selectively thinned by a second DRIE step to the intended shank thicknesses of 15, 25 and 50 µm using a photo resist etch mask. The silicon based untethered test structures are similarly fabricated, but we omitted the metallization and electrode coating steps. In contrast, the tethered silicon-based test structures were realized in identical manner as the recording probes and bonded to a short PI cable without an interface to the external instrumentation. The cables connected to the silicon based neural probes were fabricated as described for the polyimide probes. Furthermore, the contact and bonding pads interfacing with a ZIF connector and the pads on the probe base were thickened with gold electroplating (Suppl. Figure 2B). We connected the probes to the cables with flip-chip bonding (Suppl. Figure 2D), as has been described in Kisban et al. (2009). Finally, the silicon based insertion shuttles (Suppl. Figure 1E) were fabricated with the EBAG process (Sharma et al., 2023) applied to single-side polished silicon wafers. Table 1 provides an overview of the probe types implanted in the different groups of mice.

**Table 1.**
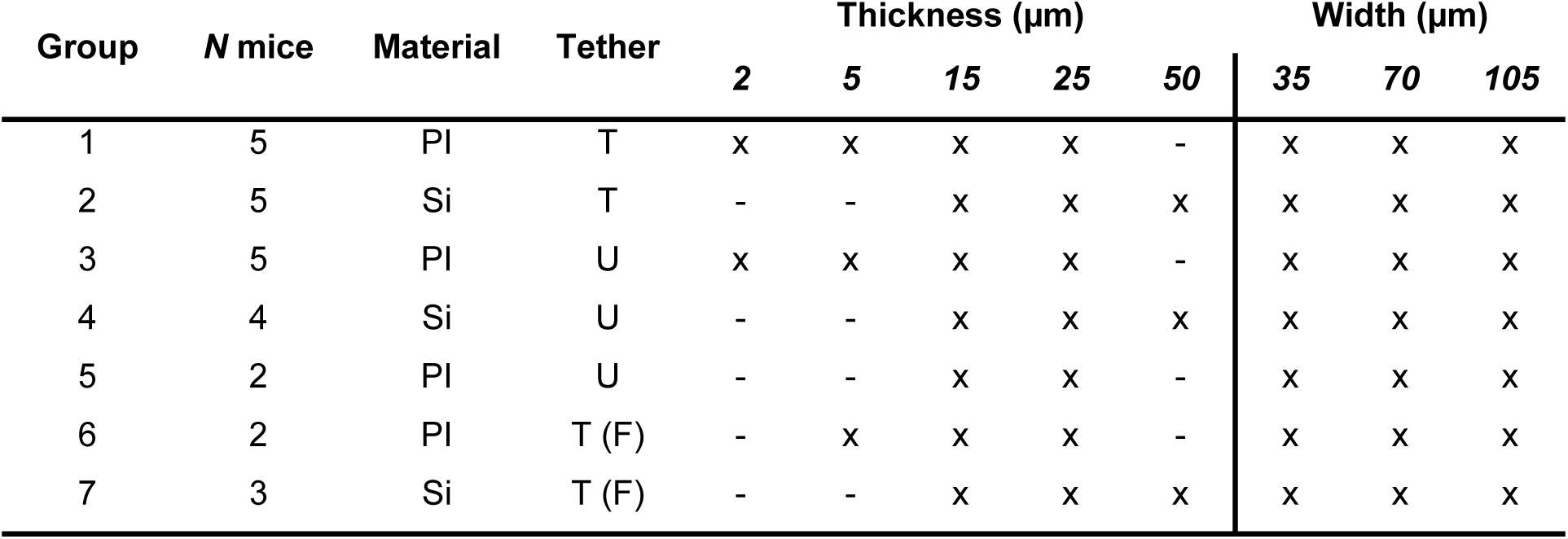
Overview of experimental groups and the implanted probes. ***PI***, Polyimide; ***Si,*** Silicon; ***T***, Tethered implantation; ***U***, Untethered implantation; ***F***, Functional electrode array. ***X***, Probe of this dimension has been implanted; ***-***, no probe of this dimension has been implanted. The number of mice reflects the mice included in our analysis and is smaller than the total number implanted mice.

### Surgical procedure

We anesthetized the mice with 4-5% isoflurane in an induction box. Anesthesia was kept constant during surgery with 1.5-2% isoflurane in an oxygen-enriched air (50% air and 50% O_2_) mixture and was monitored throughout the surgery. For analgesia we used a subcutaneous injection of 5 mg/kg Metacam and we also injected 8 mg/kg dexamethasone intraperitoneally. We used a heating pad to maintain a body temperature between 36.5 and 37.5 °C.

We fixed the head of the mouse in a stereotaxic set-up and applied ocular cream on the eyes to prevent eye dehydration. We shaved and cleaned the head of the animal and applied Xylocaine on top of the head prior to the first incision for additional local analgesia. We cut a single vertical incision in the skin along the midline and exposed the skull between Bregma and Lambda. We cleaned the skull and applied dental primer to ensure fixation of cement.

The surgical implantation procedure differed between tethered vs untethered probes. Tethered probe combs were attached to the skull, whereas untethered electrode combs could move with the brain. To implant tethered probes, we drilled horizontal cranial slits of 1.5 mm in the bone above both hemispheres. The probes were glued to a brass bar which served as an implantation device mounted on the stereotactic arm. The three shanks were inserted at a depth of 1.4 mm. For the polyimide probes we used a silicon shuttle device. Upon probe insertion, the wings of the polyimide structure were embedded in cement that was applied to the skull. We applied saline to dissolve the PEG from the brass bar and to detach the polyimide probes from the shuttle device. We closed the cranial slits with dental cement (Figure 1B top). To implant untethered probes, we drilled a circular craniotomy of approximately 3 mm in diameter in the bone above one or two hemispheres, exposing the underlying cortex. The implantation device was retracted upon implantation and we fixed a stainless-steel cap above the craniotomy with dental cement, closing the craniotomy. This cap had a 660 μm inner roof to prevent pressure on the implanted probes. (Figure 1B bottom).

In mice with non-functional probes, we stitched up the skin and sterilized the wound. For mice implanted with functional electrode arrays, we drilled an additional craniotomy over the right or left frontal cortex to place a ground wire under the skull that made contact with the cerebrospinal fluid. We placed the PCB at a vertical angle on the mouse skull to allow for access to the ZIF-connector and attached the PCB, the wiring and a head-bar to fixation the mouse with dental cement.

At the end of the implantation period (6-12 months) we euthanized the mice by deeply anesthetizing them with an intraperitoneal injection of 0.3 ml pentobarbital (60 mg/ml) and perfused them with phosphate buffered saline (PBS) followed by 4% paraformaldehyde (PFA). We extracted the brains and temporarily stored them in PFA for post-fixation for 24 hours. The following day we transferred them to 30% sucrose.

### Lesion identification and cryosectioning

We dyed the brains for 20 minutes in 4% Evans Blue, a solution which stains for albumin. When the brain blood barrier is disrupted, albumin permeates brain tissue, where it is otherwise absent, and can be visualized by Evans Blue binding. We capitalize on this reaction for the identification of lesions created by the surgical insertion of the probes.

We marked an area of 1.5 mm above and 1.5 mm below the insertion sites of probes as regions of interest, on the dorsal surface of the intact brains. Once marked, the brains were snap frozen in isopentane at -50°C and stored at -80°C. We cut 50 μm-thick, free-floating coronal cryosections of the region of interest for immunohistochemistry, collected in 1:1 Glycerol:PBS and stored at -20°C. To remove any glycerol excess, we washed the free-floating cryosections in PBS for five minutes. We then stained the sections in 4% Evans Blue for 20 minutes, for further visualization of the lesion sites within each section. We identified and assigned lesions to the respective probe types and only selected sections with clear insertion sites for further analysis. On average, this included 4 sections per probe lesion. We stored sections in PBS with 0.1% sodium azide (NaN_3_), to prevent any microbial contamination when stored at 4°C.

### Immunohistochemistry

We washed selected sections in PBS for five minutes prior to the staining protocol. We then blocked and permeabilised the sections with 2% normal donkey serum (NDS; Jesser et al., 2023) in Permeant Solution (PS) (Lai et al., 2023) for one hour. PS is composed of 0.3% TritonX and 0.1% sodium azide in PBS, which allows for eased tissue permeabilisation and prevents microbial contamination. Following blocking, the sections were stained first for 10 minutes with primary antibodies; mouse-NeuN (MA5-33103 Invitrogen, 1:500), goat-GFAP (SAB2500462 Sigma, 1:500) and rabbit-IBA-1 (019-19741 Fujifilm, 1:1000) in PS with 5% NDS at 37°C and thereafter overnight at room temperature (Biran et al., 2005). We used these primary antibodies to stain mature neurons, reactive astrocytes and microglia, respectively (Table 2). The sections were washed in PBS for five minutes and then stained with secondary antibodies at a final concentration of 1:500 in PBS (Table 3). We kept sections in the dark to prevent photobleaching. We first dyed sections for ten seconds with Alexa 647 at a higher concentration (3:500 in PBS), and immediately after we added Cy3 and Alexa 594 to reset the concentration of the secondary antibodies at 1:500. Staining Alexa 647 at a higher concentration for those initial 10 seconds ensured IBA-1 staining to not be overshadowed by NeuN and GFAP. The sections were first placed for ten minutes on a moving plate ensuring homogenous staining and then incubated at 37°C for two hours. We then washed the sections for five minutes at room temperature in PBS and mounted them on SuperFrost slides (Fisher Scientific, Epredia, ref. J1800AMNZ) with antifade mounting medium Vectashield (Vector Labororatories, ref. H-1000), which preserves the fluorescence signal and prevents rapid photobleaching. The slides were stored at 4°C in the dark and were ready for microscopic analysis.

**Table 2.** Primary antibodies (AB).

| Primary AB | Manufacturer | Concentration | Host |
| --- | --- | --- | --- |
| NeuN | Invitrogen | 1:500 | Mouse |
| GFAP | Sigma | 1:500 | Goat |
| IBA-1 | Fujifilm | 1:1000 | Rabbit |

**Table 3.** Secondary antibodies (AB).

| Secondary AB | Manufacturer | Concentration | Host | Reactive Species | Emission Peak [nm] | Excitation Peak [nm] |
| --- | --- | --- | --- | --- | --- | --- |
| Cy3 | Jackson<br>ImmunoResearch | 1:1000 | Donkey | Mouse | 555 | 569 |
| Alexa 594 | Jackson<br>ImmunoResearch | 1:1000 | Donkey | Goat | 590 | 618 |
| Alexa 647 | Jackson<br>ImmunoResearch | 1:1000 | Donkey | Rabbit | 650 | 671 |

### Image preprocessing

In total we imaged 252 stained sections with a Leica SP8 confocal laser scanning microscope system at 1024 × 1024 pixel resolution and a scan speed of 100 lps. We selected sections for preprocessing and analysis that exhibited clear lesions of a probe shank in one of the ROIs (see details below). We selected 165 experimental images and 5 control images for further analysis. Control images came from the non-implanted hemispheres of 5 mice in the different experimental groups. We preprocessed images with the FIJI software (Schindelin et al., 2012). Images were oriented to position the lesion from the 35 μm-wide shank on the left side and from the 105 μm-wide shank on the right side of the image. We cut images around the individual shank lesions to a width of 800 pixels. The height of the images was determined by biological markers. We placed the top of the image at the upper edge of the pial surface of the cortex. In images in which full layers of cortex were missing due to the damage caused by the probe, we inferred the position of the upper edge based on the full image of the section. We determined the lower end of the image as 700 pixels below the boundary of cortex and the underlying corpus callosum. In images in which probes were too posterior and no underlying white matter was present we determined the lower edge as the end of the cortical hemisphere wing.

### Channel Signal Quantification

We used two segmentation techniques to identify the signal of the stained cells. First, we threshold the images using the automated threshold method *Moments* developed by Tsai (1985) which categorizes the individual channel images into a binarized output of cell vs non-cell. We then cut the binary shank specific images into 3 depth regions of interest, each of a size of 800 × 700 pixels (∼461 × 404 μm): 1) upper cortex, 2) lower cortex, and 3) white matter. We obtained a total of 447 NeuN, 451 GFAP, and 383 IBA1 ROI images (Table 4). As a second segmentation step, we built a mask of the image by filtering the binarized image. The filtering criteria varied for the three stains based on the different morphologies of neurons, astrocytes and microglia. This allowed us to clean the image of any light artefacts and background noise. As a final step we applied a smoothing factor by down sampling the ROI images to a resolution of 11 × 10 pixels (Figure 1C,D).

**Table 4.**
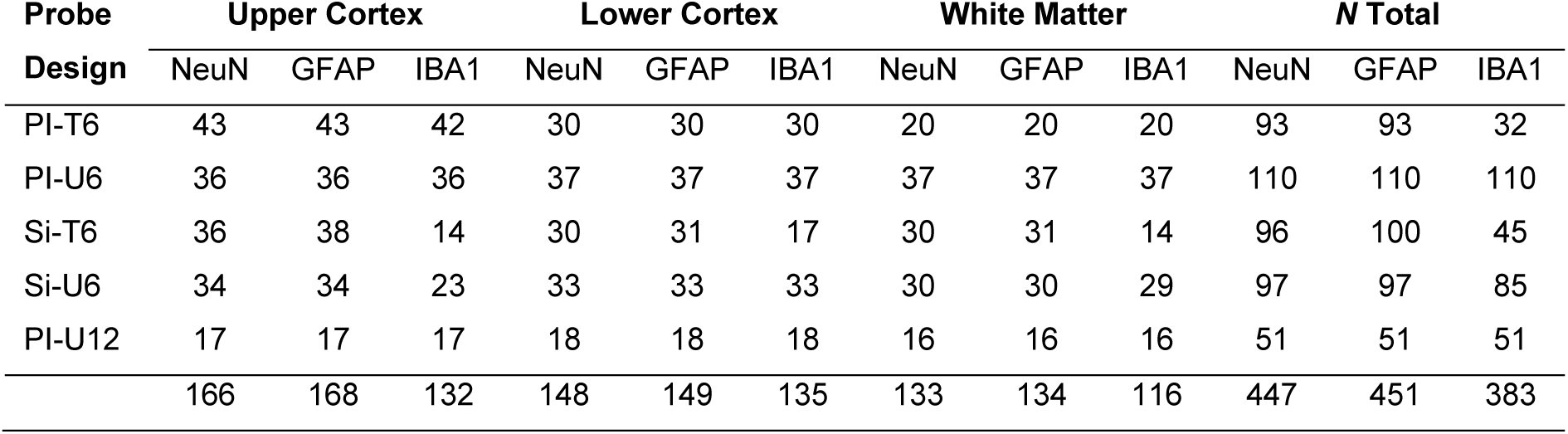
Number of sample images per probe design and staining marker. Sample numbers are presented per ROI and total number. ***PI***, Polyimide; ***Si,*** Silicon; ***T***, Tethered implantation; ***U***, Untethered implantation; ***6***, 6-month implantation duration; ***12***, 12-month implantation duration.

| Probe<br>Design | Upper Cortex |  |  | Lower Cortex |  |  | White Matter |  |  | N Total |  |  |
| --- | --- | --- | --- | --- | --- | --- | --- | --- | --- | --- | --- | --- |
|  | NeuN | GFAP | IBA1 | NeuN | GFAP | IBA1 | NeuN | GFAP | IBA1 | NeuN | GFAP | IBA1 |
| PI-T6 | 43 | 43 | 42 | 30 | 30 | 30 | 20 | 20 | 20 | 93 | 93 | 32 |
| PI-U6 | 36 | 36 | 36 | 37 | 37 | 37 | 37 | 37 | 37 | 110 | 110 | 110 |
| Si-T6 | 36 | 38 | 14 | 30 | 31 | 17 | 30 | 31 | 14 | 96 | 100 | 45 |
| Si-U6 | 34 | 34 | 23 | 33 | 33 | 33 | 30 | 30 | 29 | 97 | 97 | 85 |
| PI-U12 | 17 | 17 | 17 | 18 | 18 | 18 | 16 | 16 | 16 | 51 | 51 | 51 |
|  | 166 | 168 | 132 | 148 | 149 | 135 | 133 | 134 | 116 | 447 | 451 | 383 |

### Tissue Loss Quantification

In many cases the probe implantation caused a loss of cortical tissue. To quantify the extent of the damage we added the NeuN and GFAP signals of the individual shank images as total tissue signal. We squared the signal, highlighting true tissue signal and suppressing background noise. We then applied a threshold of <50 signal intensity, creating a binarized categorization of tissue vs non-tissue. We inverted the image to define tissue loss with a high signal intensity and tissue with low intensity, which were analyzed per depth region and down-sampled to 11 × 10 pixels (Figure 1E). The complete dataset of tissue loss can be found in Suppl. Figure 4A.

### Signal analysis

We analyzed the histological data by calculating the difference in signal between the experimental and control condition by subtraction. For analyses in which we focused on the complete depth profile of the image we calculated the delta values on a pixel-by-pixel basis. For group and condition analyses we calculated the average over pixels per depth area before calculating the delta values. To account for tissue loss in the analyses of channel signals, we removed pixels displaying >75% tissue loss (>190 tissue loss signal intensity) from the individual experimental images. For the NeuN analyses we additionally removed the first 3 rows of the upper cortex ROI in both experimental and control conditions, which represented layer 1 of the cortex with few neurons (Figure 2E, control image). Averages are calculated as means over sample images and presented with the standard error. The complete datasets of ΔNeuN, ΔGFAP and ΔIBA1 can be found in Suppl. Figure 4B,C.

To examine total tissue reaction, defined by neuronal loss and immune increase, we developed an index, Idx_TissueReaction_, that takes the ΔNeuN and ΔGFAP signal into account. We did not include ΔIBA1 because we only had a few images in the tethered silicon group in which the stain took hold. Per sample image we added the absolute value of the ΔNeuN signal to the ΔGFAP signal. Probe insertion will generally not cause neuronal genesis or astrocytic loss so that positive ΔNeuN values and negative ΔGFAP values are likely cause by biological variation. We therefore set those signal values to zero prior to the channel addition. We normalised Idx_TissueReaction_ by dividing the signal by 510 because the signal in each channel ranges from 0 to 255 so that the index ranges between 0 (no tissue reaction) and 1 (complete neuronal loss and strong astrocytic reaction).

### Statistical analysis

The statistical analysis of ΔNeuN was restricted to the cortical ROIs, because there are few neurons in the white matter. Group differences in ΔNeuN were first assessed using analysis of variance (ANOVA). Significant main or interaction effects were followed up with post-hoc or pairwise t-tests, applying Bonferroni correction to account for multiple comparisons. In contrast, ΔGFAP and ΔIBA1 values did not meet the assumptions of normality and were therefore analyzed using non-parametric statistics. Group effects were assessed using Kruskal–Wallis tests, followed by post-hoc Dunn tests or pairwise Dunn tests with Bonferroni correction where appropriate. Sample numbers presented with the statistical test results are based on the total number of cut ROI images included in the calculation. Table 4 gives an overview of the number of sample images per probe designs and staining markers.

### Electrophysiology

We performed electrophysiological recordings of 5 mice during passive viewing of a visual stimulus, for 24 weeks post-implantation. Mice were head-fixed on a running-wheel in front of a monitor. Visual stimuli consisted of a full screen checkerboard that was presented for 500 ms, after which the pattern inverted (black checks became white and vice versa) and was present for another 500 ms. The mice saw 100 stimuli with an inter-trial interval of 1s. Electrical signals were amplified and sampled at 24 kHz using a Tucker Davis Technologies recording system. We re-referenced the signal to the common average to remove running and muscle artefacts. Multi-unit-activity (MUA) responses were detected by band-pass filtering the signal between 750-5000 Hz, rectifying it by taking the absolute, and applying an additional low pass filter of 200 Hz (Supèr & Roelfsema, 2005). We calculated the signal to noise ratio (SNR) by smoothing the MUA responses and defined noise as the standard deviation over trials of the mean signal 250 ms prior to stimulus onset. The visual response magnitude was measured as maximum absolute value of the mean signal deviation from the baseline in a time-window from 0–250 ms (checker-board presentation), 500–750 ms (switch), or 1–1.25 s (stimulus off) post stimulus onset, and we chose the response with highest magnitude. In the calculation, we used the absolute value of the signal because we also observed phases of response suppression below baseline, representing response inhibition. In weeks with multiple sessions, we averaged the SNRs across sessions to obtain a single SNR value per week. We analyzed signal quality over time by fitting a linear regression model to the SNR of electrodes of polyimide and silicon probes, and we also computed the percentage of useful electrodes (SNR > 1) for each material. Differences between polyimide and silicon probes over time were calculated by analyzing the SNR data in six time-bins of four weeks each with Dunn tests. We investigated the influence of probe dimension (thickness and width) with the two electrode materials with Kruskal-Wallis tests, followed by post-hoc pairwise Dunn tests with Bonferroni correction.

## Acknowledgements

We thank the staff of the animal department for biotechnical support. The work was supported by NWO grants (Crossover grant 17619 ‘‘INTENSE’’ and ‘‘DBI2,’’ a Gravitation program of the Dutch Ministry of Science), ZonMW grant 09120232310021 ‘EVISION’, LSH Match Project LSHM23002 (POSITIONED), the European Union Horizon 2020 Framework Program under grant agreement 899287 ‘‘NeuraViper,’’ an ERC grant (101052963 ‘‘NUMEROUS’’), ERC-POC grant (101138075, “PROVISO”) and RO1 grant R01EY036094 to P.R.R, as well as EBRAINS 2.0 HORIZON-INFRA-2022-SERV-B-01 (101147319) awarded to R.N.K.

## Supplementary Figures

**SUPPL. FIGURE 1.**
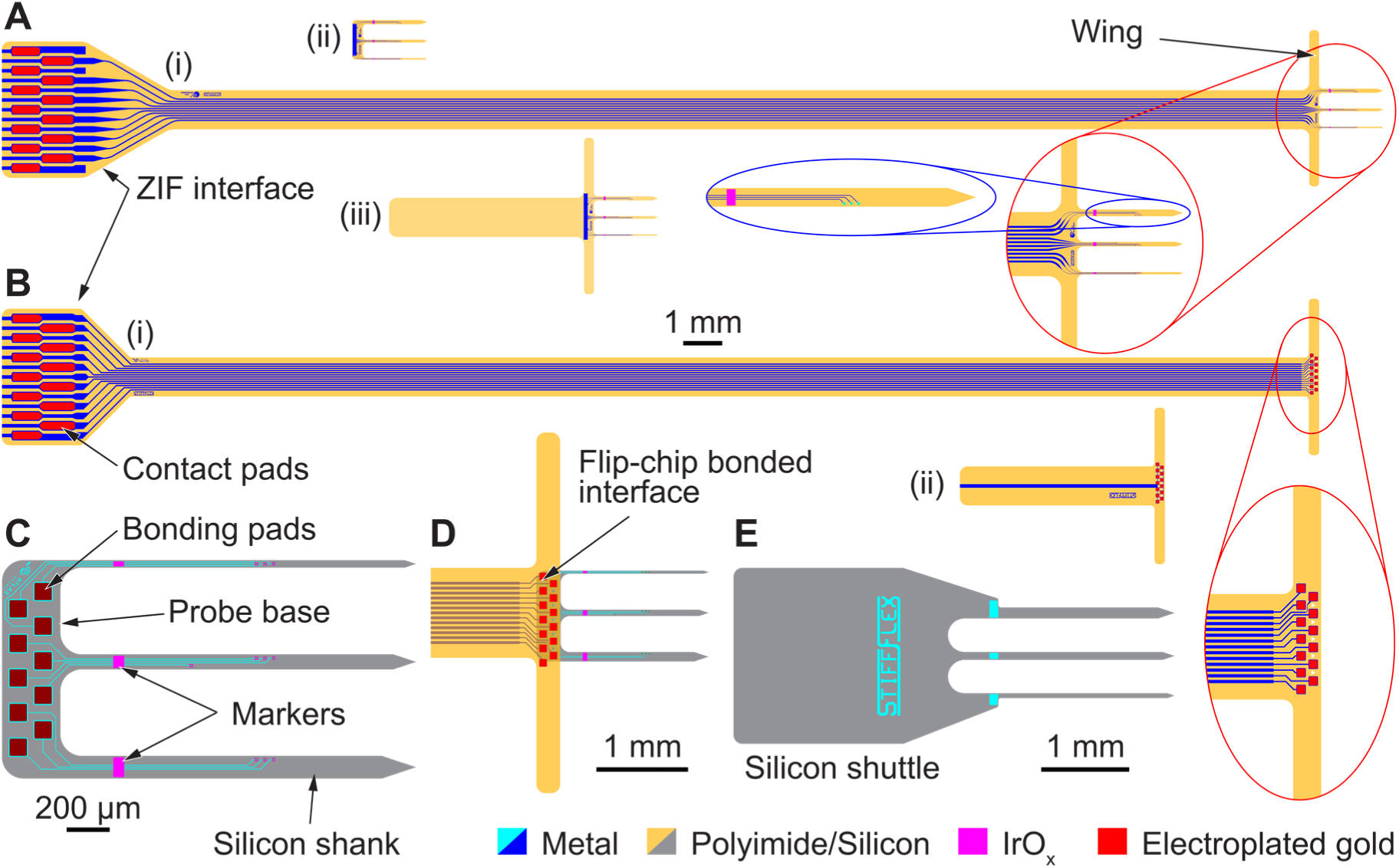
Neural probe designs. (**A)** Polyimide probes and test structures with three implantable shanks: (i) Functional probe with a 28-mm-long cable connected to a ZIF interface. The wing structure is used to fix the probe after insertion to the skull with dental cement; (ii) floating implant with a slim base interconnecting the probe shanks, and (iii) tethered implant with a short PI strip mimicking the cable in (i). **(B)** Interconnecting cables for (i) the Si-based probe used for neural recording and (ii) dummy cable with a short PI strip mimicking the cable in (i). **(C)** Silicon probe with 10 bonding pads to which the cable from (B-i) or cable dummy from (B-ii) are flip-chip bonded. **(D)** Flip-chip bonded interface between the Si-based neural probe and the PI-based cable. **(E)** Silicon-based insertion shuttle with a larger base to facilitate the handling of the probe.

**SUPPL. FIGURE 2.**
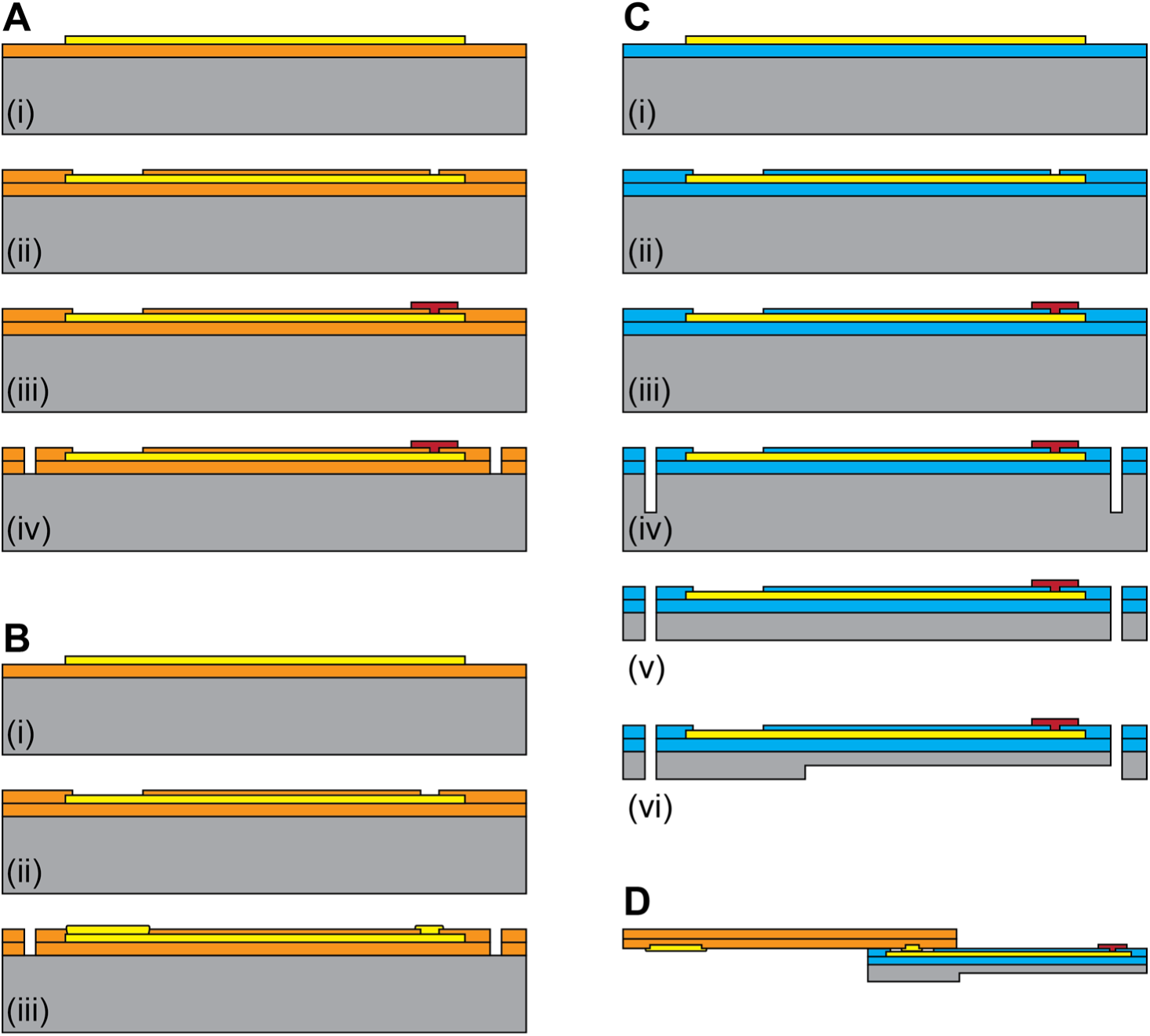
Fabrication process. (**A)** Fabrication of PI-based neural probe: (i) Spin-coating of the first polyimide layer, and deposition and patterning of metallization (100 nm Pt) using sputtering and a lift-off process, (ii) spin-coating of the second polyimide layer and RIE to open recording sites (electrodes) and contact pads, (iii) reactive sputter deposition and lift-off patterning of IrO_x_ electrode coating on top of the polyimide layer, and (iv) patterning of neural probe contour using RIE. **(B)** Fabrication of polyimide interconnecting cable: (i) Spin-coating of first polyimide layer, and deposition and patterning of metallization (100nm Pt) using sputtering and a lift-off process, (ii) spin-coating second polyimide layer and RIE to open contact and bonding pads and definition of the cable contour, and (iii) electroplating of contact and bonding pads with gold. **(C)** Fabrication of silicon probes: (i) Deposition of the first passivation layer containing a layered stack of SiO_x_ and Si_x_N_y_ using PECVD, and deposition and patterning of metallization (100 nm Pt) using sputtering and a lift-off process, (ii) deposition of second passivation layer stack of SiO_x_ and Si_x_N_y_ using PECVD and RIE patterning of recording sites and contact pads, (iii) reactive sputter deposition and lift-patterning of IrO_x_ electrode coating on top of the SiO_x_/Si_x_N_y_ passivation layer, (iv) patterning of probe contour by RIE and bulk micromachining of silicon using DRIE, (v) wafer grinding to thin down the silicon substrate, and (vi) selective DRIE to thin down the probe shanks using the EBAG process. **(D)** Silicon probe assembly: Flip-chip bonding of the silicon probe to the polyimide interconnecting cable.

**SUPPL. FIGURE 3.**
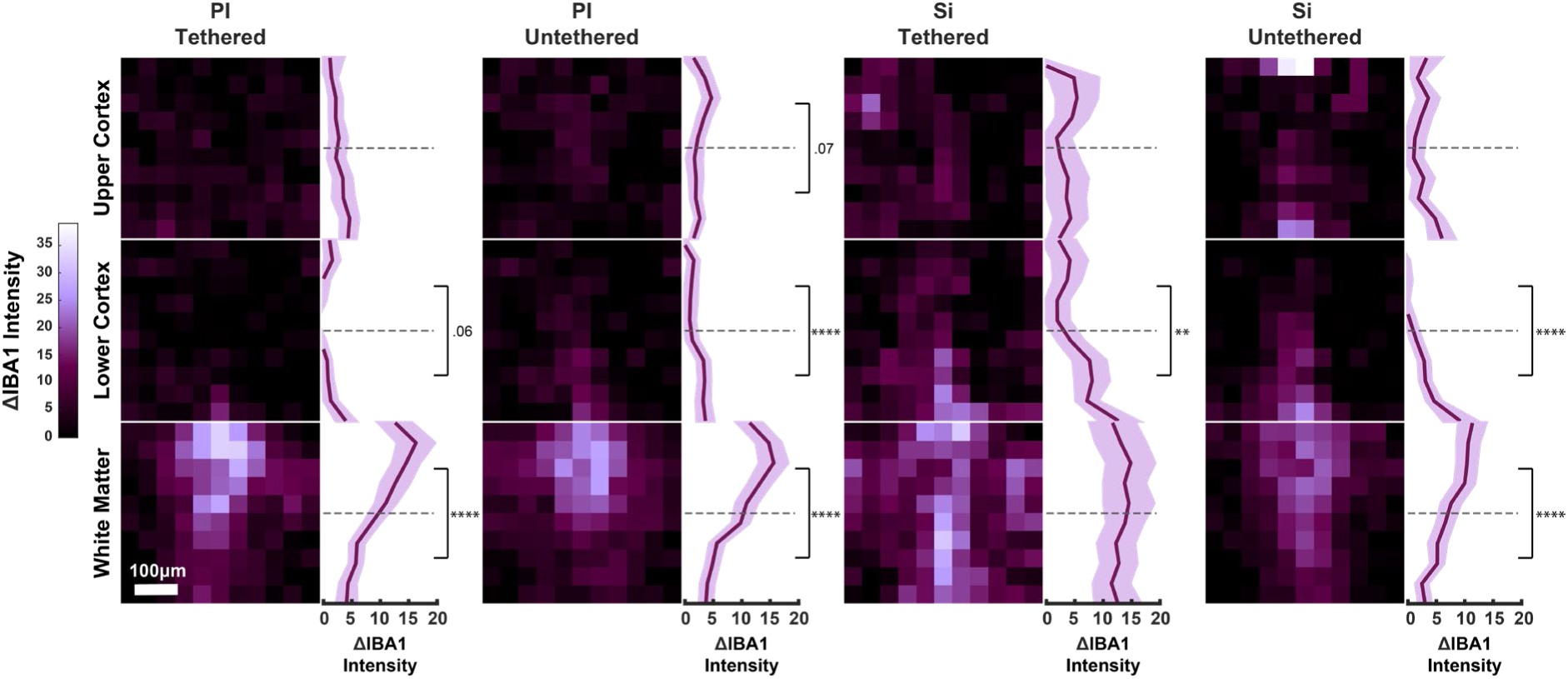
Microglial depth profile for the tethered and untethered silicon and polyimide probes. Scale bar = 100 μm. PI = Polyimide; Si = Silicon. Stacked heatmaps correspond to upper cortex (top), lower cortex (middle) and white matter (bottom). Line graphs display the average ΔIBA1 value across cortical depth. Shaded areas represent SEM. Significant differences of signal intensities in upper and lower areas within each ROI are marked. All conditions display an increased ΔIBA1 intensity in the white matter. * p < .05, ** p < .01, *** p < .001, **** p < .0001.

**SUPPL. FIGURE 4.**
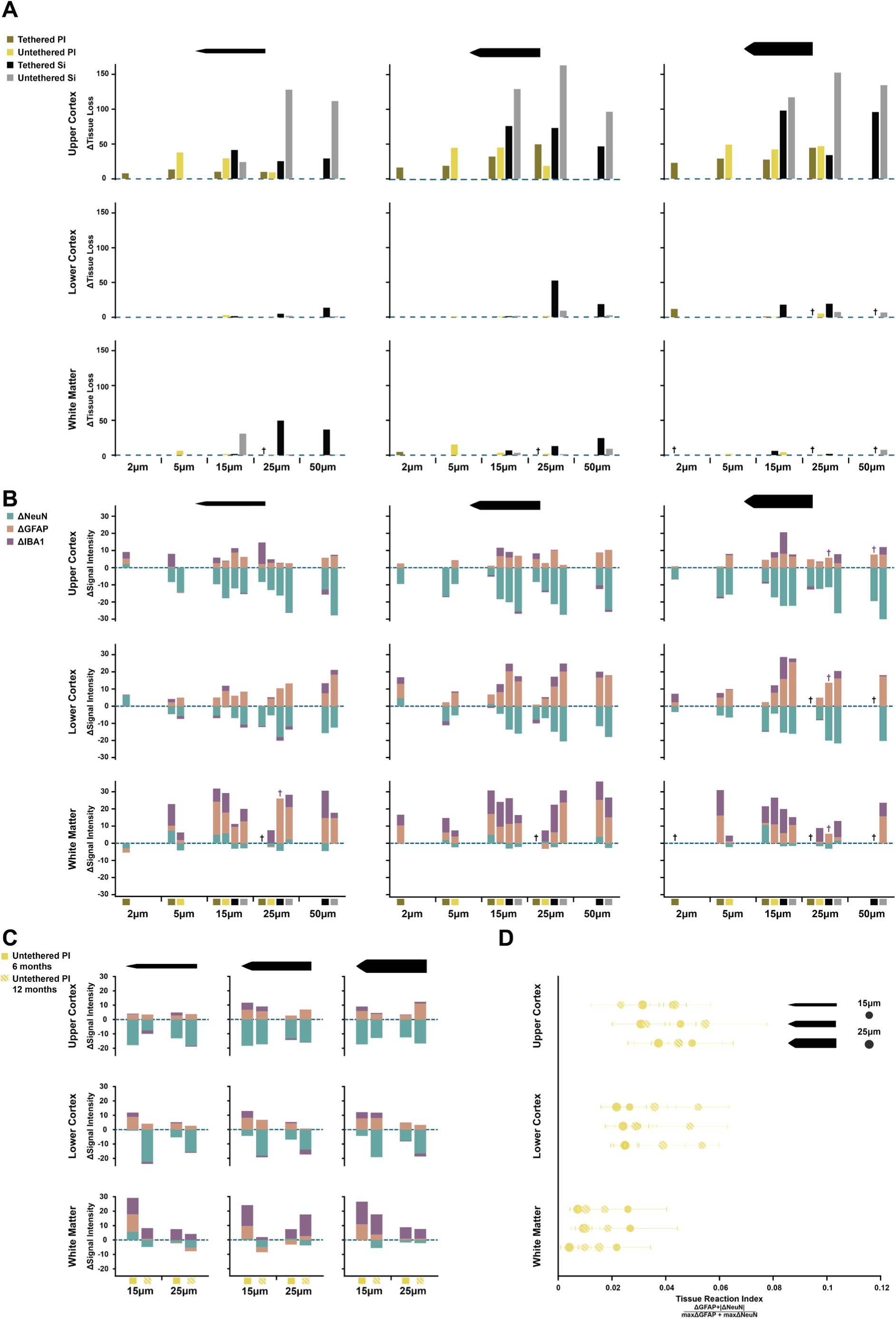
Tissue responses across all experimental conditions. **(A)** ΔTissueLoss caused by the various probes with a shank of 35 (left), 70 (middle column), 105 µm width (right) in upper cortex (top), lower cortex (middle row) and white matter (bottom). **(B)** ΔNeuN (neuronal loss), ΔGFAP (astrocytic response) and ΔIBA1 (microglial response), in the same format as panel A. The colors below the x-axis represent the various material and tethering combinations. Black plusses mark missing datapoints for conditions with one sample or less. Colored plusses mark missing datapoints in a specific staining. **(C)** Untethered polyimide probes implanted for 6 or 12 months of shanks with a width of 35 (left), 70 (middle column) and 105 µm shank width (right column). **(D)** Idx_TissueReaction_ for untethered polyimide probes implanted for 6 and 12 months.

**SUPPL. FIGURE 5.**
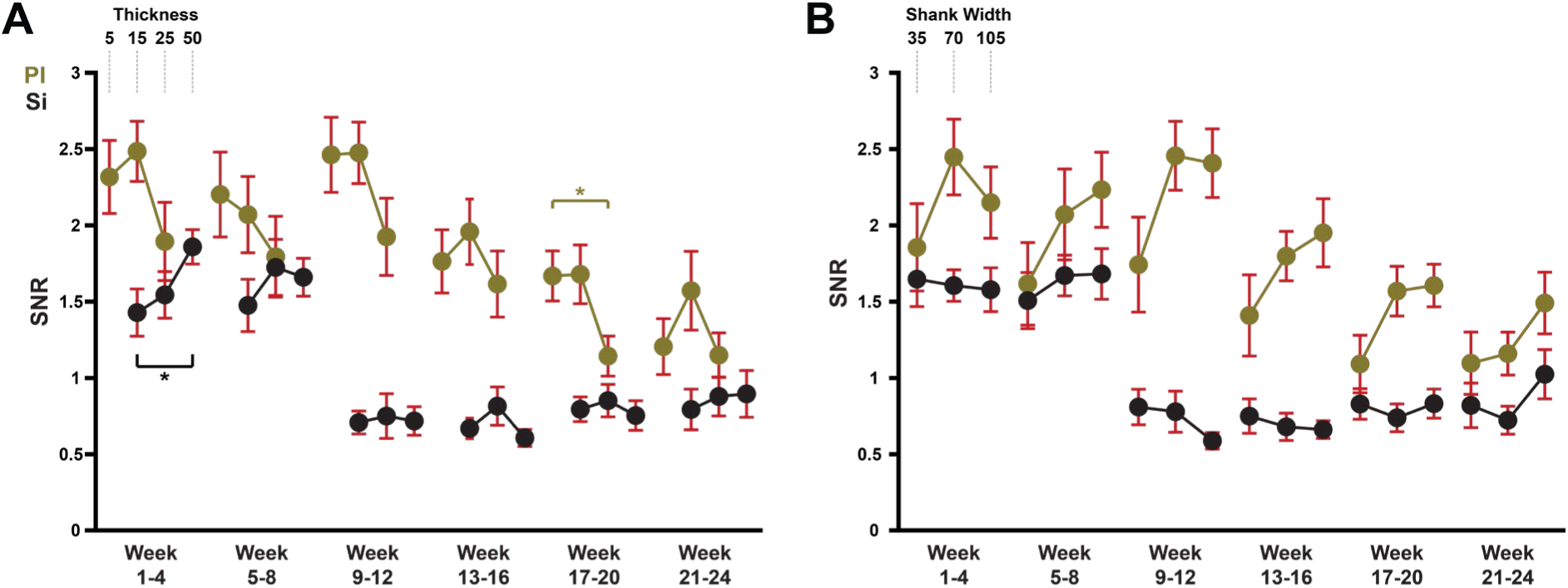
Average signal-to-noise (SNR) ratio for electrodes on shanks with different thicknesses and widths in 4 week time-bins. PI = Polyimide; Si = Silicon. Error bars, SEM. **(A)** Influence of probe thickness on SNR. Polyimide probes had a thickness of 5, 15, or 25 µm and silicon probes were 15, 25 or 50 µm thick. **(B)** SNR of electrodes from shanks with a width of 35, 70 or 105 µm. * p < .05.

**SUPPL. FIGURE 6.**
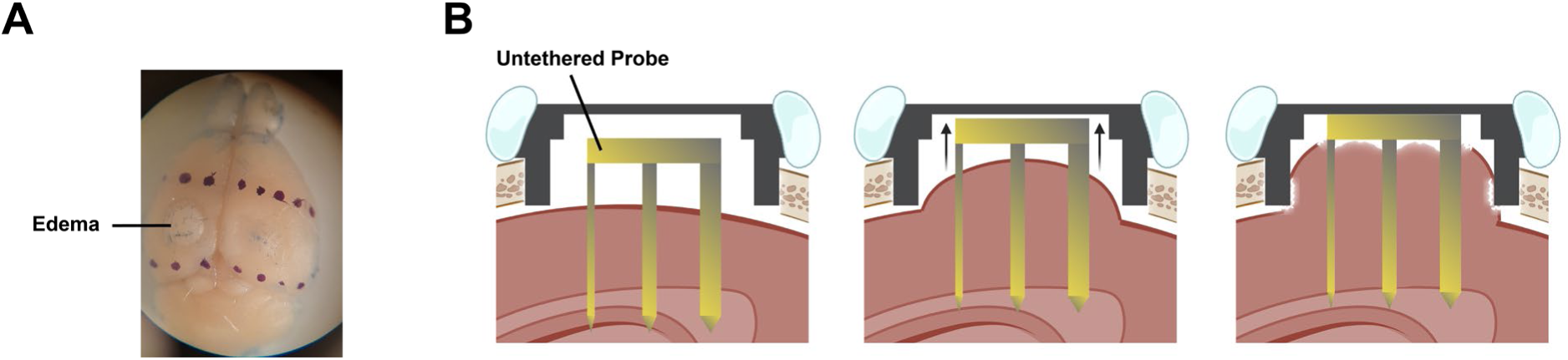
Adverse effects of the surgical method used to implant untethered probes. We made a 3 mm circular craniotomy and closed the skull with a stainless-steel cap. **(A)** Example brain with edema at the site of the implantation in the left hemisphere. **(B)** Illustration of the possible negative impact of the cavity above the cortex. Pressure may cause displacement of the cortex and probe into the cavity in the direction of the inner roof of the cap.

